# The Role of Polyglutamine in Inter- and Intra-molecular Interactions in Med15-dependent Regulation

**DOI:** 10.1101/2022.12.29.522246

**Authors:** David G. Cooper, Jan S. Fassler

## Abstract

Med15 is a general transcriptional regulator and member of the tail module of the RNA Pol II Mediator complex. The *S. cerevisiae* Med15 protein has a well-structured N-terminal KIX domain, three Activator Binding Domains (ABDs) and several naturally variable polyglutamine (poly-Q) tracts (Q1, Q2, Q3) embedded in an intrinsically disordered central region, and a C-terminal Mediator Association Domain (MAD). We investigated how the presence of ABDs and changes in length and composition of poly-Q tracts influences Med15 activity and function using phenotypic, gene expression, and transcription factor interaction assays of truncation, deletion, and synthetic alleles. We found that individual Med15 activities were influenced by the number of activator binding domains (ABDs) and adjacent polyglutamine composition. We also observed that distant glutamine tracts and Med15 phosphorylation affected the activities of the KIX domain, suggesting that intramolecular interactions may be required for KIX domain interactions with transcription factors. We conclude that robust Med15 activity required at least the Q1 tract and that the length of that tract modulates activity in a context-dependent manner. We speculate that the glutamine tract provides a degree of intramolecular flexibility that is needed for Med15 function. Finally, we found that loss of Msn2-dependent transcriptional activation in Med15 Q1 tract variants correlates well with a reduction in Msn2:Med15 interaction strength.

## Introduction

Polyglutamine (poly-Q) tracts, or consecutive repeats of the amino acid glutamine, are a dynamic protein feature with the potential to cause disease by expansion of the underlying microsatellite repeats [1-3]. However, poly-Q tracts have been ascribed non-pathogenic roles that include providing flexible intradomain spacing and protein-protein interaction surfaces [4-8]. Perhaps not surprisingly many of the proteins containing poly-Q tracts are transcriptional regulators [9]. Polyglutamine tract length differences encoded by naturally occurring alleles of genes have been shown to modulate protein activity. Variation in the length of the poly-Q tract of the Clock protein corresponds to alterations in barn swallow breeding [10]. The subcellular localization of Angustifolia in *Populus tremula* is altered by poly-Q tract length changes [11]. Variation in the length of the poly-Q tract of the Notch protein alters developmental phenotypes in fruit flies [12]. Natural and synthetic Q and QA repeat variability in yeast Cyc8 has broad transcriptomic effects [13]. We hypothesize that similarly variable poly-Q tracts in yeast Med15 alter the transcriptional activation function of the protein.

Poly-Q tracts could influence activity by affecting protein-protein interactions, either directly or indirectly; providing disorder to allow a larger set of transient interactions; providing necessary spacing between functional domains; or by providing flexibility to the protein. Poly-Q proteins have a higher number of interactions than other proteins and poly-Q proteins are often found in protein complexes [9]. One interaction surface mediated by poly-Q tracts are coiled-coil domains [14]. In yeast, homodimerization of the Nab3 component of the Nrd1-Nab3 transcription termination complex is facilitated by the coiled-coil structure of a C-terminal Q-tract [5, 15]. The polyglutamine content in the human Med15 ortholog has been shown to form a coiled-coil structure [16]. However, how the multiple poly-Q tracts in yeast Med15 may influence transcriptional activation has not been thoroughly explored.

As a subunit of the RNA Polymerase II Mediator complex, Med15 functions as an interaction hub with other transcriptional regulators to regulate gene expression. Distributed throughout the Med15 sequence are residues and domains that have been shown to correspond to the interactions between Med15 and other proteins. The C-terminal Mediator Association Domain (MAD (aa 799-1081)) permits Med15 to assemble into the Mediator complex [17, 18]. Specifically, conserved amino acid residues 866-910 in MAD are required for Med15 incorporation into Mediator [19, 20]. The domains or regions of Med15 required for interactions with yeast transcription factors (TFs) including Oaf1, Pdr1/3, Msn2, Gcn4, and Gal4 have been mapped with varying resolution [21]. The N-terminal KIX domain (aa 6-90) mediates interactions between Med15 and the Pdr1/3 and Oaf1 TFs [22, 23]. The interactions with other TFs require sequences within the glutamine-rich central region of Med15 and are discussed below.

Med15, formerly Gal11, was initially discovered in conjunction with its role in galactose metabolism [19, 24]. Med15 regulates the expression of galactose metabolism genes through interactions with Gal4 bound to UAS_G_ motifs upstream of *GAL* genes [25]. Another well-studied Med15 interactor is Gcn4. Gcn4 regulates many genes in yeast, including those involved in amino acid biosynthesis [26, 27]. The Gcn4 TF interacts with residues in 4 regions of Med15, the N-terminal KIX domain and three so-called Activator Binding Domains (ABD1 (aa 158-238), ABD2 (aa 277-368), and ABD3 (aa 496-630)) embedded in the intrinsically disordered and glutamine-rich midsection of the protein [20]. Gcn4 and Gal4 interactions with Med15 are described as “fuzzy” meaning that the interactions can be mediated using different subsets of the available hydrophobic residues in interaction pockets [28, 29]. Interacting Gcn4 and Med15 proteins form liquid droplets [30]. Liquid droplet formation is one form of phase separation and serves as a mechanism for concentrating the two interacting proteins. Liquid droplet phase separation may be one way to increase the concentration of interactors to compensate for “transient” interactions.

Med15 also interacts with the core environmental stress response transcription factors Msn2/4 to regulate gene expression of stress response genes [31]. Med15 regulates the expression of Msn2 dependent genes *HSP12* and *TFS1* [31]. Although Med15 is not known to interact directly with Hsf1, expression of some heat shock proteins including *SSA4* and *HSP104* is also regulated by *MED15*.

The central section of Med15 (∼aa 100-700) is enriched in amidic amino acids glutamine (16% of residues) and asparagine (11% of residues) [21]. Within this region there are three variable polyglutamine (poly-Q) tracts (longer than 10 residues) which we refer to as Q1 (aa 147-158), Q2 (aa 417-480), and Q3 (aa 674-696) that flank, but are not included in the ABD regions. Q1 and Q3 are simple poly-Q tracts while Q2 consists of alternating glutamine-alanine repeats. While shorter stretches of glutamine are present in Med15, Q1, Q2 and Q3 stand out because they vary in length (number of consecutive residues) in *MED15* alleles from different strains of *S. cerevisiae* [21]. Recent studies have addressed the functional implications of naturally occurring tract length variant alleles of *MED15* on resistance to the coal-cleaning chemical, 4-methylcyclohexane methanol [32], and grape juice fermentation [33]. Here we investigate which additional activities of Med15 require these glutamine-rich regions, and the extent to which variations to the polyglutamine tracts in Med15 influence these protein activities.

## Results

The ability of yeast to respond to stress and changes in growth conditions requires the expression of target genes dependent on interactions between Med15 and gene-specific transcription factors. To test the extent to which distinct transcription factor interacting regions and the intervening sequences in Med15 contribute to its activity, we analyzed a series of internal deletions and truncated *MED15* alleles (Fig. 1A) for phenotypes exhibited by the deletion mutant (Fig. 1B). All *MED15* alleles are driven by the native *MED15* promoter.

**Figure 1.**
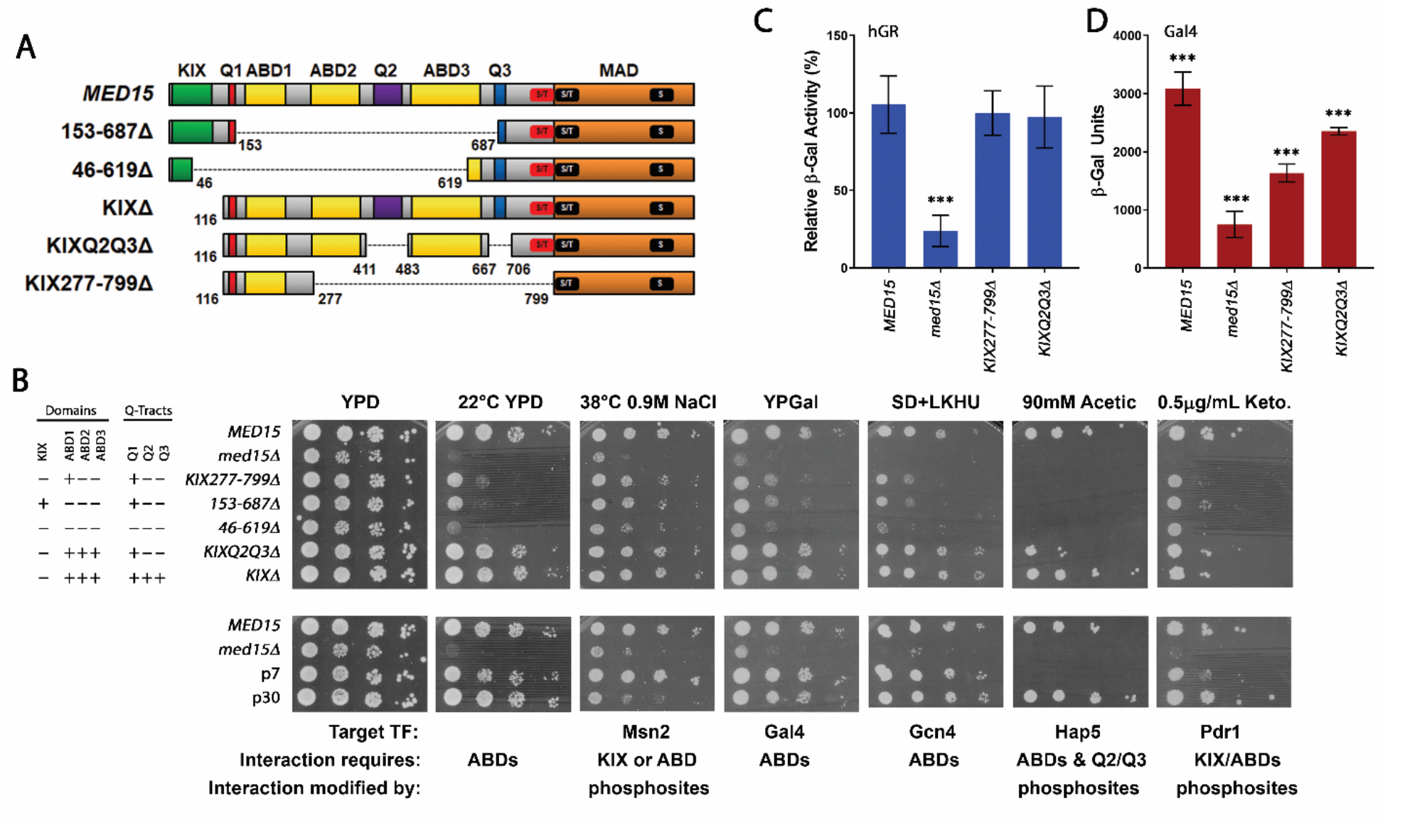
Internal deletions and associated Med15 activities. **(A)** Full length Med15 (1081 aa) domains: KIX (Kinase-inducible domain Interacting Domain), ABD1-3 (Activator Binding Domains), Q1-3 (polyglutamine tracts), MAD (Mediator Association Domain). Regions containing phosphorylation sites (S/T and S) including those dynamically phosphorylated in response to salt stress (red box) and others (black boxes). Numbers represent amino acid residues. KIXΔ-*MED15* constructs lack the KIX domain. KIXQ2Q3Δ-*MED15* constructs lack the KIX domain and the poly-Q tracts for Q2 and Q3. In addition to the construct with the lab allele 12 glutamine residues at Q1, KIXQ2Q3Δ-*MED15* constructs include a 0-Q1 construct as well as versions with different Q1 tract lengths (#-Q1) and non-glutamine sequences. KIX277-799Δ-*MED15* constructs lack the KIX domain and central Q2/Q3 containing region. **(B)** Log-phase cultures of the *med15Δ* strain (OY320) carrying a CEN plasmid with the indicated *MED15* allele, or strains with integrated phospho-site mutations (D7P and D30P) were serially diluted and spotted on YPD media with or without supplements and incubated at 30°C, 38°C, or room temperature (22°C). **(C-D)** β-galactosidase activity of a human glucocorticoid receptor dependent (hGRτ1-*lacZ)* reporter (C, blue) or Gal4-dependent (UAS_G_-*lacZ*) reporter (D, red). **(C)** β-galactosidase activity of a human glucocorticoid receptor dependent (hGRτ1-*lacZ*) reporter in *med15Δ* strain (JF1368) carrying a copper inducible hGRτ1-LexA plasmid, an episomal LexA-*lacZ* reporter, and a CEN plasmid with the indicated *MED15* allele. **(D)** β-galactosidase activity of a Gal4-dependent (UAS_G_-*lacZ*) reporter in *med15Δ gal80Δ* strain (JF2631) carrying a CEN plasmid with the indicated *MED15* allele. Data points are the averages of at least 3 biological replicates (transformants) and error bars are the standard deviation of the mean. Significant differences were determined using ANOVA analysis with a Tukey post-hoc test, *** p≤0.001.

### Four Med15 Activator Binding Domains contribute additively to Med15 activity

Two internal deletion constructs (153-687Δ and 46-619Δ) lacking most of the central region of Med15 displayed different levels of activity when expressed in a *med15Δ* strain (Fig. 1). Deletion 46-619Δ disrupts or lacks all characterized activation domains, while deletion 153-687Δ still maintains an intact KIX domain (Fig. 1). Both constructs contain the Mediator Association Domain (MAD), which is necessary for Med15 activity [17, 19]. The 46-619Δ construct failed to complement most of the tested *med15Δ* defects and others it complemented very weakly (Fig. 1B). Maintenance of the KIX domain, as in the 153-687Δ construct, was sufficient for partial to full complementation of the tested phenotypes (Fig. 1B).

### The ABD1 Region Alone (Q1R) Partially Complements *med15Δ* Defects

KIX277-799Δ-*MED15* retains the glutamine-rich region of Med15 around Q1 (Q1R; aa 116-277) and the MAD domain (aa 799-1081) (Fig. 1A) but lacks the KIX domain, ABD2, ABD3, Q2 and Q3. When expressed in a *med15Δ* strain, KIX277-799Δ-*MED15* partially complemented many of the *med15Δ* phenotypes tested, including utilization of galactose as a carbon source and tolerance to ethanol (not shown) (Fig. 1B), and provided nearly full complementation for growth in the presence of combined osmotic and thermal stress. As previously reported, the KIX277-799Δ-*MED15* construct also fully activated an hGR-dependent reporter gene (Fig. 1C) in the presence of a partial (mammalian) hGR transcription factor [18].

Only one of the two regions of the Med15 protein involved in interactions with the Gal4 transcription factor (aa 116-176 and aa 566-618) [34] is included in the KIX277-799Δ-*MED15* construct. The presence of this one region (aa116-176) allows for partial activation of a Gal4-dependent reporter (Fig. 1D). The reduced level of expression relative to full length *MED15* is consistent with the phenotype of KIX277-799Δ-*MED15* on galactose plates (Fig. 1B).

### KIX domain activity

Specific activities have been attributed to the discretely-folded and well-conserved KIX domain (aa 6-90), including interactions with the transcription factor Oaf1 to regulate the expression of fatty acid metabolism genes [23] and interactions with Pdr1/3 to regulate the expression of drug efflux pumps involved in pleiotropic drug resistance [22, 35]. The KIX domain is also known to contribute to Gcn4 and Gal4 activation alongside other activator binding domains (Fig. 1A, ABD1-3 annotations) [20, 28, 36, 37].

Here we use ketoconazole tolerance as a phenotypic indicator of Pdr activity. Constructs lacking the KIX domain exhibited mild sensitivity to the drug, however a construct containing the KIX domain but lacking other ABDs was also mildly sensitive, suggesting the involvement of other Med15 regions in activation of the Pdr response (Fig. 1B). In contrast, the presence or absence of the KIX domain alone (compare KIXΔ to *MED15*) appeared to have little detectable impact on Msn2 (heat, osmotic stress) or Gal4 (growth on galactose as sole carbon source) and Gcn4 (growth on SD media) phenotypes. The importance of the KIX domain in combination with ABD1-3 for Gal4 dependent activation could also be seen in the KIX domain-containing *med15* 153-687Δ mutant, which exhibits better growth on galactose plates compared to the *med15* 46-619Δ mutant, which lacks an intact KIX domain. In constructs lacking the KIX domain, the ABDs are important. For example, KIX277-799Δ -*MED15* and the KIXQ2Q3Δ−*MED15*, both lack the entire KIX domain but contain either a single (ABD1; KIX277-799Δ) or all three ABDs (KIXQ2Q3Δ) and exhibit intermediate (KIX277-799Δ) or excellent (KIXQ2Q3Δ) growth on galactose (Fig. 1B), although reporter assays indicate that both constructs have lower expression of Gal genes than the wild type allele (Fig. 1D). Similarly, growth on SD media, a reflection of Gcn4 activity, is poor in the 46-619Δ mutant in which both the KIX domain and the ABD domains are partially or entirely absent and best in the KIXQ2Q3Δ mutant which lacks the KIX domain but retains ABD1, ABD2 and ABD3 (Fig. 1B).

### The Central Q-rich and Intrinsically Disordered Region

We simplified the analyses of the central disordered domain and its three long Q tracts by analyzing them in the absence of the KIX domain. The KIXQ2Q3Δ-*MED15* construct which lacks the KIX domain and the Q2 and Q3 glutamine tracts, fully complements all tested phenotypes except for ketoconazole and acetic acid tolerance when expressed in a *med15Δ* background (Fig. 1B) suggesting that the region around Q1 is more important than Q2 and Q3 for most KIX domain-independent Med15 functions. Consistent with this conclusion, the KIXQ2Q3Δ-*MED15* construct fully activated the hGR-dependent reporter (Fig. 1C). Interestingly, acetic acid tolerance, a likely function of the recently uncovered interaction between Med15 and Hap5 [38] was sensitive to the absence of Q2 and Q3 (Fig. 1B).

### Med15 Phosphorylation

In unstressed cells Med15 is phosphorylated, a modification proposed to dampen Med15 activation of stress genes. In contrast in cells exposed to osmotic challenge Med15 is dephosphorylated [39]. There are 30 phosphorylated residues in Med15, all clustered around the junction of the central disordered domain (distal to Q3) and MAD (Fig. 1A) [39]. We examined two mutants to begin to assess the effect of phosphorylation on Med15 activity in response to other stresses. In the D7P mutant, 7 residues whose phosphorylation is affected by osmotic stress were changed to alanine, and in the D30P mutant, all 30 phospho-sites were changed to alanine. A weak growth defect at high salt concentrations was previously reported for the D30P mutant [39]. In contrast the D7P mutant was not affected by high salt. We found that these mutants differed in response to acetic acid as well. D30P exhibited wild type tolerance to acetic acid stress, while D7P was highly sensitive to acetic acid stress and was less tolerant to ketoconazole (Fig. 1B).

### The Q1 Tract and Length variants

KIXQ2Q3Δ-*MED15* constructs containing a normal size Q1 tract consisting of 12 glutamines (12Q) fully complemented several of the tested phenotypes including the utilization of galactose as a carbon source; sensitivity to ethanol; and sensitivity to NaCl at 37°C (Fig. 1, Fig. 2A). However, a construct lacking the Q1 tract (0-Q1 KIXQ2Q3Δ-*MED15*) neither fully complemented *med15Δ* defects on plates (Fig. 2A), nor reporter assays of hGR and UAS_G_-*lacZ* activation (Fig. 2C, D). Additional resolution was achieved by quantifying growth in liquid media containing a stressor versus growth in media with no addition. In general, 0Q and 6Q inserts at Q1 afforded better tolerance than the *med15* deletion strain, but much less than 12Q. In some cases, 36Q and 47Q constructs exceeded 12Q (Fig. 2B: acetic acid, ketoconazole) and in other cases 36Q and 47Q were less robust than 12Q (Fig. 2B: ethanol).

**Figure 2.**
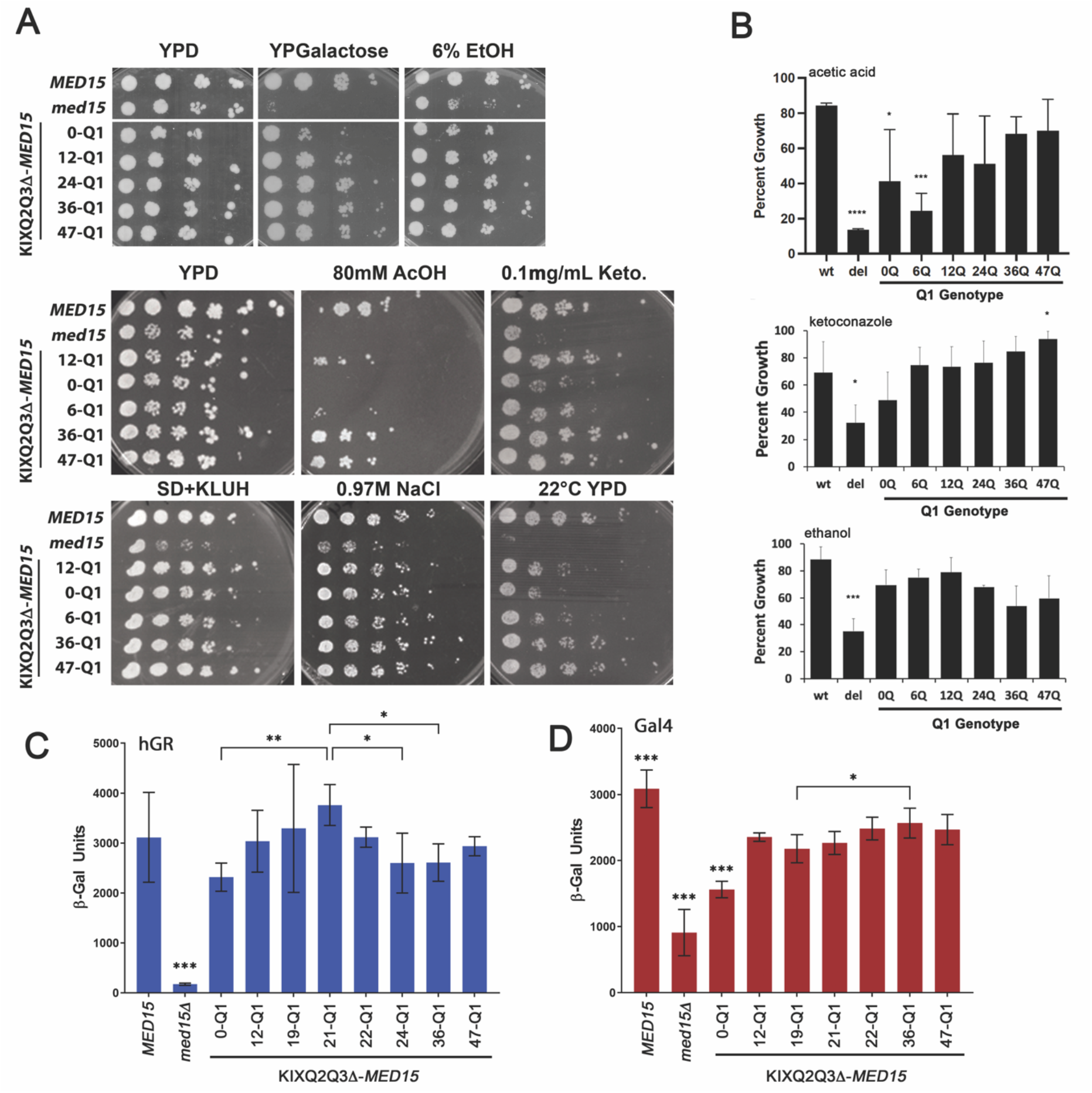
Poly-Q length variation at Q1 alters Med15 activity. **(A)** Log-phase cultures of the *med15Δ* strain (OY320) carrying a CEN plasmid with the indicated *MED15* or KIXQ2Q3Δ-*MED15* allele were serially diluted and spotted on YPD media with or without supplements and incubated at 30°C or room temperature (22°C). **(B)** Quantitative microtiter dish assay for growth in various media or in the presence of added chemicals. Each genotype was tested in biological triplicate. Plotted values are averages of the endpoint OD_600_ divided by growth in permissive conditions (media without chemical or YPD, 30°C). Drug concentration was chosen for an effect on the wild type MED15 strain of no more than 50% reduction in growth. **(C-D)** β-galactosidase activity of a human glucocorticoid receptor dependent (hGRτ1-*lacZ)* reporter (C, blue) or Gal4-dependent (UAS_G_-*lacZ*) reporter (D, red) as in Figure 1. All data points are the averages of at least 3 biological replicates (transformants) and error bars are the standard deviation of the mean. Significant differences were determined using ANOVA analysis with a Tukey post-hoc test, * p≤0.05.

Tract length effects at Q1 were context dependent. A Q1 tract length of 21 glutamines (Fig. 2C) exhibited a significant increase in hGR activity over constructs having 24 or 36 glutamines at Q1. In contrast, changes to the Q1 tract length had only a minor impact on Med15 activation of a Gal4-dependent reporter (Fig. 2D). These results suggest that while the Q1 tract is not essential for the activities tested, it is required for full Med15 activity and different Med15 activities are differentially sensitive to length at Q1.

### Med15 Activity Depends on the Chemical Nature and Coiled-Coil Propensity of the Q1 Tract

The region around the Q1 tract is predicted to form coiled-coil structure (Fig. 3A and Supplementary Fig. 1) which could serve as a site for protein-protein interactions. A series of modified Q-tracts and non-Q insertions into the Q1 position of the KIXQ2Q3Δ construct were analyzed to test the coiled-coil sequence requirements at the Q1 locus (Fig. 3A). A glycine-rich spacer (spacer: PGSAGSAAGG), was used to test whether a flexible sequence without coiled-coil propensity would perturb Med15 activity. The spacer construct was comparable to 12-Q1 under conditions in which the absence of the Q1 tract impaired Med15 activity (Fig. 3B, spacer). The Q1 locus was also replaced with a glutamine tract with dispersed proline residues (12PQ and doubled, 24PQ: PQQQPQQPQQQP), a sequence previously shown to disrupt coiled-coil structure [14] and predicted to reduce the coiled-coil structure of the adjacent region in Med15 (Fig. 3A and Supplementary Fig. 1). The proline interrupted sequences displayed activity comparable to 12-Q1 (Fig. 3B). In contrast, the introduction of leucine residues (15LQ; LQQQLQQLQQQLLLQ) previously shown to promote coiled-coil structure [14] did not complement the defects exhibited by the 0-Q1 construct (Fig. 3B top, most evident on galactose media).

**Figure 3.**
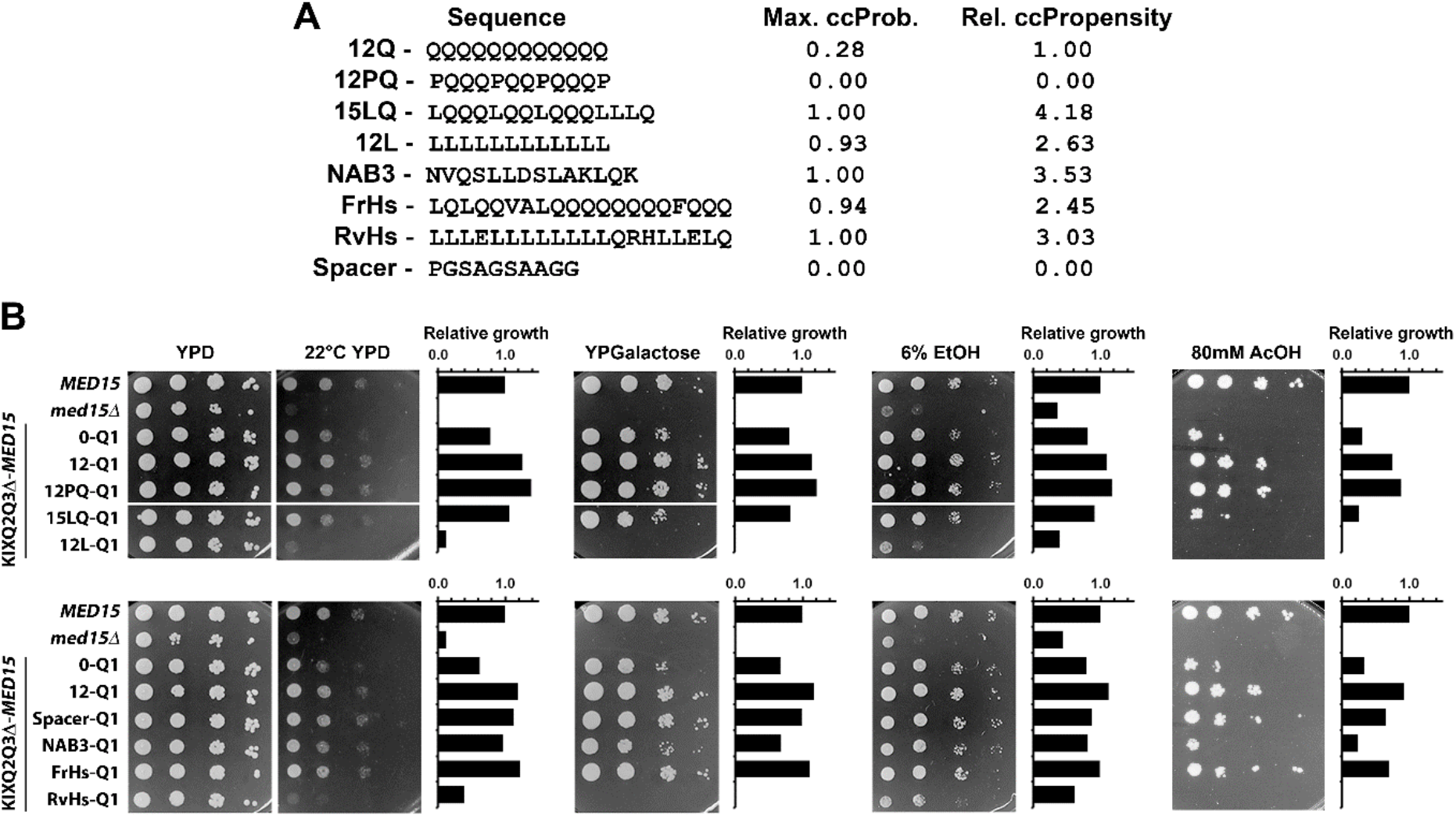
Q1 tract region coiled-coil structure perturbs Med15 activity. **(A)** Sequence and coiled coil (cc) propensity of non-Q inserts. Maximum ccProbability (Max ccProb.) corresponds to the peak coiled-coil formation probability within the Q1 region for each construct. Relative ccPropensity (Rel. ccPropensity) is the average ccProbability across the Q1 region peak relative to the 12Q construct prediction. **(B)** Log-phase cultures of the *med15Δ* strain (OY320) carrying a CEN plasmid with the indicated *MED15* allele were serially diluted and spotted on YPD media with or without supplements and incubated at 30°C or room temperature (22°C). A pixel-based relative growth measurement that incorporates both viability and colony size) was determined per plate relative to the wild type *MED15*^+^ strain.

To further investigate the apparent deleterious effect of coil-coil promoting sequences, several additional coiled-coil forming structures [5, 16] were introduced. The first pair was designed to encode a 20 amino acid segment of the coiled coil-forming human MED15 Q-tract sequence (FrHs: LQLQQVALQQQQQQQQFQQQ) and the leucine rich sequence encoded by the reverse complement (RvHs: LLLELLLLLLLLQRHLLELQ). The second was a C-terminal helix forming sequence from the *NAB3* gene (NAB3: NVQSLLDSLAKLQK). Like 15LQ-Q1, NAB3 failed to fully complement the defects of the 0-Q1 construct (Fig. 3B middle: most evident on acetic acid containing media). Replacement by the human *MED15* Q-tract coding sequence on the other hand was roughly comparable to 12-Q1 (Fig. 3B middle: FrHs-Q1). Poly-L (12L) and the reverse complement of the human Med15 Q tract (RvHs-Q1) (a periodically interrupted poly-L tract) were both generally nonfunctional (Fig. 3B).

The effect of length and composition of the Q1 tract on target gene expression was quantified by q-rtPCR in log-phase cultures (rich media, 30°C) of strains expressing wild type *MED15* or KIXQ2Q3Δ-*MED15* constructs with various Q1 genotypes. Basal expression of selected *MED15* target genes was reduced in the absence of *MED15* (*med15*Δ) (Supplementary Fig. 2). We found that basal expression was also dependent on the presence of the Q1 tract and that Q1 tract length variants gave distinctive patterns of expression. Med15 without a Q1 tract (Q1-0) had less activity than Med15 with a Q1 tract of 12 for genes such as *AHP1* (activation) and *MET10* (repression) (Fig. 4A). Although the basal expression of *MET10* and *AHP1* was dependent on the Q1 tract, it was independent of tract length. In contrast, *SSA1, HSP12*, and *GLK1* were sensitive to tract length, peaking when the Q1 tract length was 24 (Fig. 4A).

**Figure 4.**
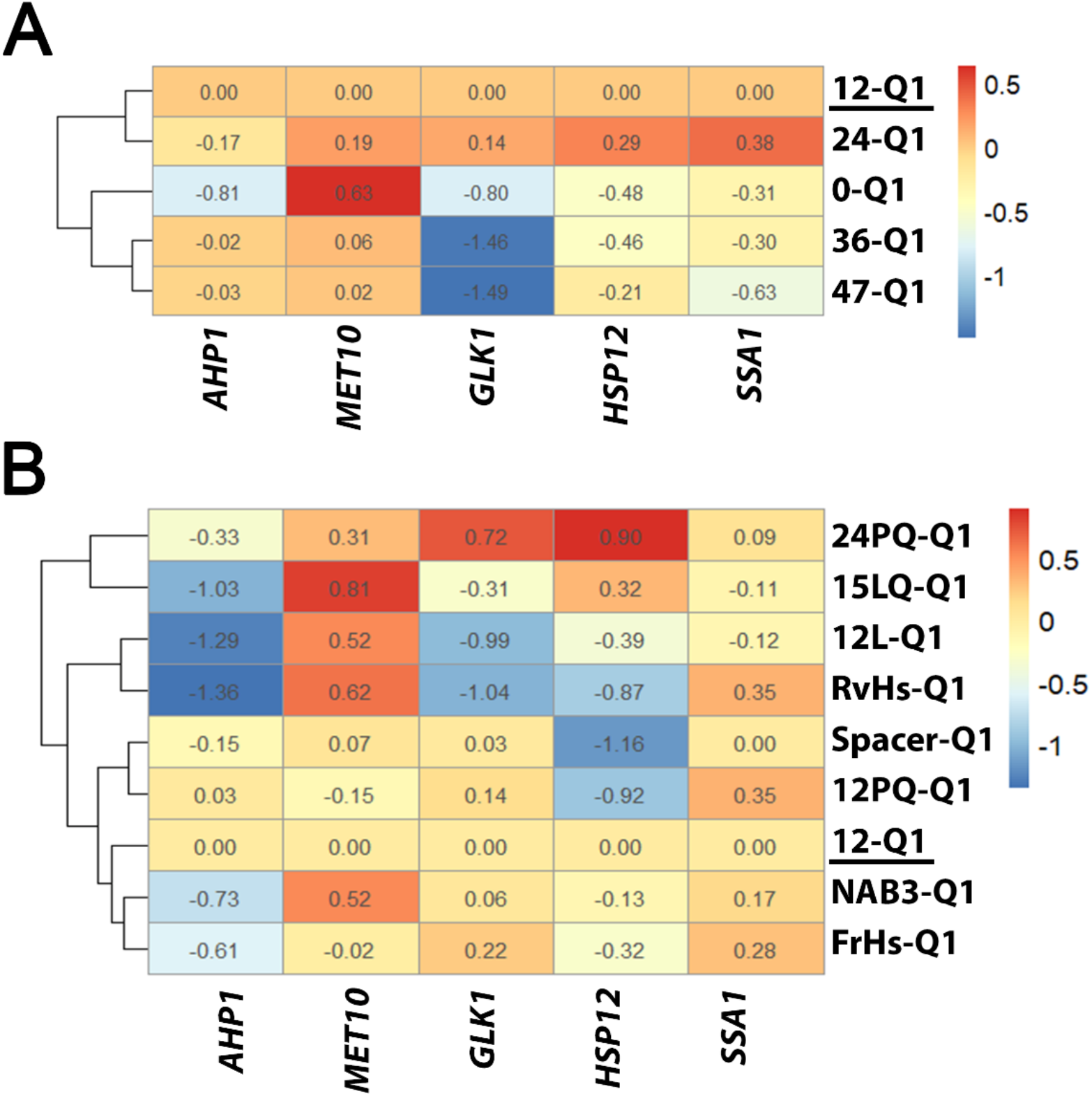
Basal Gcn4 and Msn2 dependent gene expression is modulated by the presence and length of the Q1 tract in Med15. **(A & B)** Log2 transformed relative expression of target genes for log-phase cultures of the *med15Δ* strain (OY320) carrying a CEN plasmid with specified KIXQ2Q3Δ-*MED15* Q1 variant construct in YPD grown at 30°C. Expression values for 12Q-Q1 (underlined) were set to 1 prior to log2 transformation. **(A)** Q1 length variants; **(B)** Q1 composition variants. For each strain the target gene expression was normalized to *ALG9* and averaged over 3 biological replicates (transformants). Heatmaps and hierarchical clusters prepared using pheatmap and hclust packages in R.

We also examined gene expression in the constructs with Q1 substitutions (Fig. 4B). We found that impaired gene expression correlated with phenotypes for constructs which failed to complement defects of the 0-Q1 constructs, including 12L-Q1 and RvHs-Q1 (Fig. 4B). For genes whose expression was most consistently influenced by the presence of the Q1 tract (*AHP1* and *MET10*) coiled-coil-promoting inserts including FrHs-Q1, NAB3-Q1, 15LQ-Q1, and 12L-Q1 displayed expression patterns similar to 0-Q1, while in most cases the expression patterns of spacer and PQ constructs were similar to 12-Q1. Overall, Q1 tract composition had a differential effect on the expression of different genes, consistent with the phenotype studies in Fig. 3.

### Med15 interacts with Msn2 via the KIX domain or the Q1R region

To determine whether Q1-dependent changes in the activity of Med15 reflect altered interactions with specific transcription factors, we examined Med15-Msn2 interactions using a split-ubiquitin two hybrid assay with a Ura3 reporter [40, 41]. Msn2 amino acids 1-271 (TAD) were fused to the C terminus of ubiquitin and an N-end rule sensitive derivative of the URA3 gene (RUra3), while parts of Med15 were fused to the N terminus of ubiquitin (Fig. 5A). The presence of interacting N-Ub and C-Ub fusions cause strains to become Ura^-^ due to the ubiquitin-mediated degradation of the Ura3 protein. The Ura-phenotype was tested in three ways: (1) with synthetic media lacking uracil (SD + K (required amino acid, lysine)); and (2) with synthetic complete media lacking uracil (SC-HLUM; see methods) on which strains harboring plasmids expressing fragments of Msn2 and Med15 that do interact would fail to grow, and (3) using media containing 5-fluoroorotic acid (5-FOA), an inhibitor of the Ura3 enzyme, on which only strains expressing interacting Med15 and Msn2 peptides would survive. Both the KIX domain (aa 1-118) alone (NUb-K), the Q1R fragment (aa 120-277) alone (NUb-Q1R-12Q) as well as the entire region (1-277) (NUb-KQ) were positive (Ura^-^ and 5-FOA^R^) for an interaction with the transactivation domain (TAD) of Msn2 (Fig. 5B). Consistent with the phenotypic analysis shown in Fig. 3, a NUb-Q1R with *NAB3* sequence at Q1 is slightly less interaction positive (more Ura^+^) than Q1R itself (best seen on SC-HLMU in Fig. 5B), while a Q1R with RvHs or 12L at Q1 only weakly interacts with Msn2 (best seen on 5-FOA in Fig. 5B) despite being positive for protein expression (Supplementary Fig. 3). A composite of interaction rank on all plate types splits the tested construct roughly into 3 groups (Fig. 5C). The worst interactors RvHs and 12L rank alongside negative controls. The NAB3-Q1 construct was the only tested representative of the coiled-coil-promoting constructs, and it is in a group of its own as a worse interactor than all but the non-functional constructs. The best interactor was the Spacer construct which slightly outperforms the remaining tested constructs. Hence the Q1 substitution phenotypes reflect the ability of Med15 to interact with Msn2.

**Figure 5.**
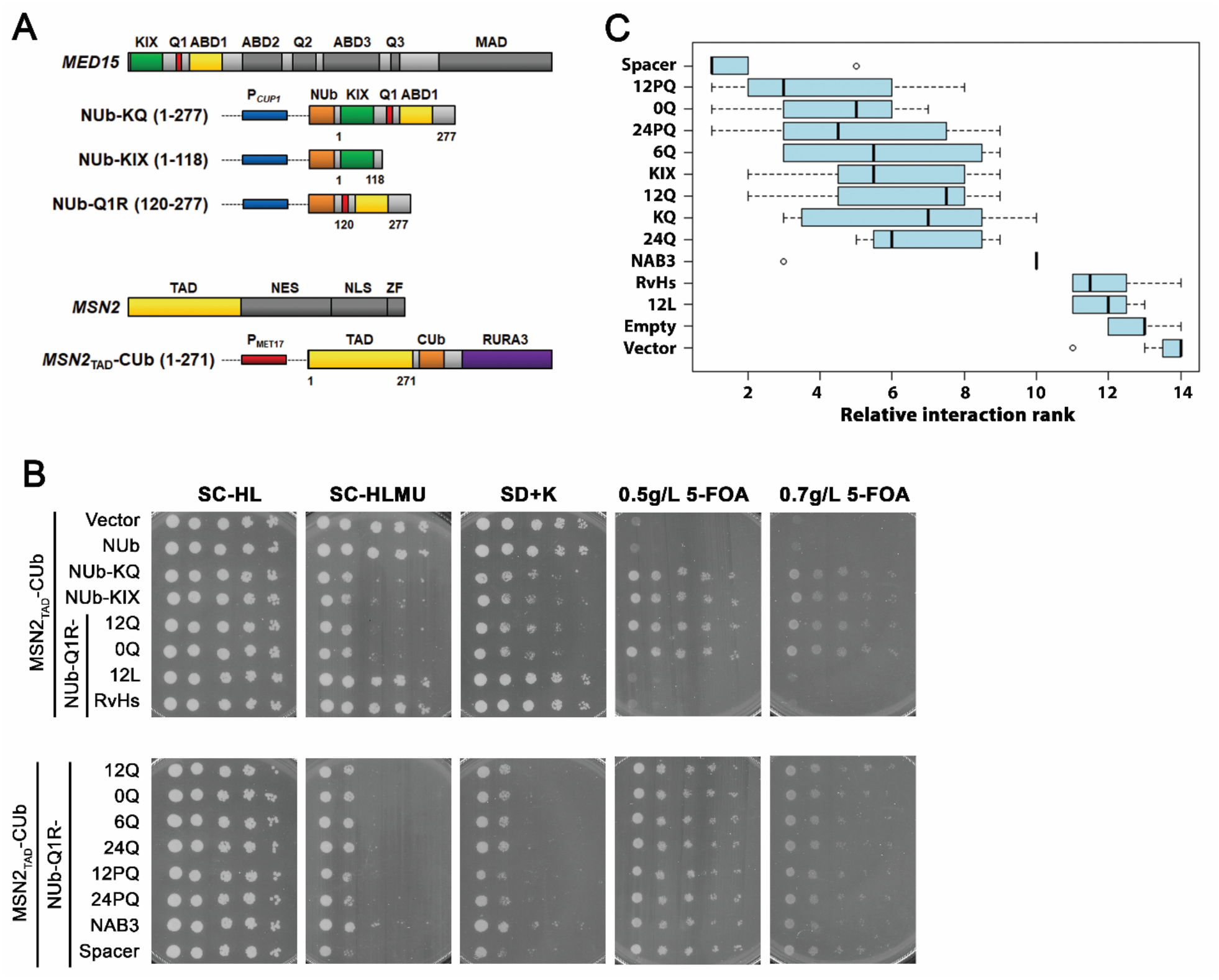
Med15 interacts with Msn2 through the KIX domain and Q1R. **(A)** Split ubiquitin constructs. Wild type yeast (BY4742) transformed with plasmids expressing N-terminal Ubiquitin fragment (NUb) fused to fragments of *MED15* behind a copper inducible *CUP1* promoter and C-terminal Ubiquitin fragment (CUb) fused to a mutated URA3 gene coding for an N-terminal arginine (RURA3) and the transcriptional-activation domain of *MSN2* (*MSN2*_TAD_) behind a methionine repressible *MET17* promoter. **(B)** Log-phase cultures in minimal media with leucine, lysine, and uracil (lacking methionine) were exposed to 0.1 or 0.2mM copper for 1 hour and then serially diluted and spotted on media lacking uracil or containing 5-FOA. Plates corresponding to strains induced with 0.2mM copper are shown. **(C)** Relative spot size formation quantified per plate as in Figure 3 at a single time point for plates lacking uracil or containing 5-FOA corresponding to strains induced with 0.1 or 0.2mM copper (8 total plates per strain). The relative growth rank was determined for each plate across all 14 tested constructs. A rank of 1 corresponds to the worst growth on plates lacking uracil and best growth on plates containing 5-FOA.

## Discussion

In this study we systematically analyzed the impact of different regions of the Med15 protein, and found that the KIX domain and Q1R region of the protein are key determinants of Med15 activity (Fig. 1) although more distal parts of the protein, including Q2, Q3 and phosphorylated residues that span the boundary with the Mediator Association Domain, appear to influence the functionality of these more N-terminal regions.

Additional insight into the role of the Q1R region of Med15 was achieved by varying the length and composition of the poly-Q tract in constructs lacking the KIX domain as well as Q2 and Q3. In these analyses, we probed the activity of the protein by assaying phenotypes that represent the output of different Med15-dependent transcription factors as well as expression of target genes. The results are summarized in Table 1. We found that activity of most tested TFs, except for Gcn4 was diminished if the Q1 tract was deleted. We further showed that several TFs except for Gcn4 and Gal4 were affected by the length of the Q1 tract, with the precise effect being context dependent. Finally, we found that the amino acid composition and/or coiled-coil propensity was a factor in the activity of Med15-dependent TFs. We observed that Q1 substitutions with increased coiled-coil propensity (Supplementary Figure 1) diminished TF activity while Q1 substitutions that interfered with coiled-coil propensity had no effect on TF activity (Fig. 3, 4), suggesting that the flexibility of the sequence is an important feature. Below we discuss each of these observations in more detail.

**Table 1.** Relationship of Med15 Q1 Genotype to TF Activity

| Phenotype | TF | Target gene /<br>reporters<br>examined | Sensitivity to Q1<br>(phenotype / gene(s)) |  |  |
| --- | --- | --- | --- | --- | --- |
|  |  |  | Presence<sup>1</sup><br>(Fig. 2,3) | Length<sup>2</sup><br>(Fig. 2,4) | Composition<sup>3</sup><br>(Fig. 3,4) |
| Acetic Acid | Hap5 | Plate assay | Yes | Yes | Yes |
| NaCl @ 38°C;<br>Ethanol | Msn2;<br>Hsf1 | Plate assays | Yes | Yes | Yes |
|  |  | GLK1 | Yes | Yes | Yes |
|  |  | AHP1 | Yes | No | Yes |
|  |  | HSP12 | Yes | Yes | Yes |
|  |  | SSA1 | Yes | Yes | Yes |
| Ketoconazole | Pdr1 | Plate assay | Yes | Yes | No*<sup>4</sup> |
| Galactose | Gal4 | Plate assay | Yes | No | Yes |
|  |  | UAS<sub>G</sub> reporter | Yes | No | NT<sup>5</sup> |
| Amino Acid<br>Limitation | Gcn4 | Plate assay | No | No | No |
|  |  | MET10 | Yes | No | No |
|  |  | ARG3 | No<sup>6</sup> | Yes* | NT |
| Suboptimal<br>Temperature<br>(22°C) | ?<sup>7</sup> | Plate assay | Yes | Yes | Yes |
| Glucocorticoid<br>Response | GR | hGR reporter | Yes | Yes | NT |

<sup>1</sup>Growth or expression altered by the removal of Q1
<sup>2</sup>Growth or expression altered by changes in the length of Q1
<sup>3</sup>Growth or expression altered by changes in the composition of Q1
<sup>4</sup>Data not shown\*
<sup>5</sup>Not tested
<sup>6</sup>Data in supplement
<sup>7</sup>Target transcription factor unknown

### Msn2 TF activity localized to two regions of Med15

A miniaturized version of Med15 consisting of aa 116-277 (ABD1, Q1) and the Mediator Association Domain (KIX277-799Δ-*MED15*) [18] fully complemented some of the stress related defects of the *med15*Δ strain. The activity of this construct is consistent with, and further refines, previous work establishing that the N terminal 351 amino acids are sufficient for interactions between Med15 and the stress responsive transcription factor Msn2 [41]. We confirmed that Msn2-dependent activities of Med15 are encoded by the region containing the Q1 tract and ABD1 (aa 116-277) and found that the KIX domain alone could also mediate an interaction with Msn2 (Fig. 5). This contrasts with the Gcn4 or Gal4-dependent growth or stress responses which are the result of additive interactions with Med15 that are characterized by weak, highly dispersed, multivalent interfaces [28, 29]. While it is not yet entirely clear if the interaction with Msn2 is similarly multivalent, we have shown that either the KIX domain alone or the Q1R region alone of Med15 was sufficient with no evidence of additivity.

We also found that Med15 Q1 substitutions affected the interaction with the Msn2 transcription factor. The Msn2 interaction with Med15 Q1 was most affected by Q1 substitutions like 12L and RvHs that were the least functional and was somewhat affected by the NAB insert which exhibited some reduction in functionality (Fig. 3, Fig. 5B, C). Hence, overall, we conclude that the nature of the Q1 sequence has an impact on the interaction with the Msn2 transcription factor, while the length of the glutamine tract at Q1 may impact Med15 function differently.

### The Hap5 TF (Acetic Acid tolerance) localized to dispersed Med15 sequences

Of the *med15* mutant phenotypes we examined, acetic acid tolerance phenotype was among the most revealing. We observed acetic acid sensitivity stemming from the absence of Q2 and Q3 and in full-length *MED15* alleles with mutations in the D7P mutant which eliminates the potential for phosphorylation required to prevent activation under non-stress conditions and which become dephosphorylated upon osmotic stress (Fig. 1) [39]. Acetic acid tolerance was also affected by the absence, length, and composition of the Q1 tract (Fig. 2, Fig. 3). Taken together the Med15 requirements for acetic acid tolerance suggests dispersed activator binding sites for the Hap5 acetic acid responsive TF [38], albeit in different locations than for Gal4 and Gcn4. An alternative explanation could be a requirement for intramolecular interactions between different regions of the Med15 protein. For example, an interaction between the phosphorylated region near the mediator association domain and N-terminal sequences like the Q1 tract may squelch Med15 activity by preventing association of the subunit with the remainder of the complex.

### The presence and length of the Q1 tract affected Gcn4 and Msn2 activation

The effect of Q1 tract length on growth and stress response phenotypes and on expression of reporters and target genes was context dependent. Basal and induced expression of individual Msn2-dependent (e.g., *AHP1*) and Gcn4 dependent (e.g., *MET10*) genes was influenced by the presence/absence of the Q1 tract (Fig. 4, Sup. Fig. 2C). In all instances TF activity was reduced in the absence of the Med15 Q1 tract. This suggests a general role for the Q1 tract in modulating Med15 dependent regulation of TFs.

Whether the length of the Q1 tract influenced gene expression depended on the transcription factor. Gal4-dependent reporter gene expression as well as the expression of Gcn4 regulated *MET10* were mostly tract-length insensitive (Fig. 2C and Fig. 4A). However, expression of an hGR-dependent reporter peaked at 21Q and was reduced at 36Q (Fig. 2B).

Similar expression patterns were seen for Msn2/Hsf1 targets including *HSP12, SSA1* and *GLK1* where expression peaked at 24Q and was reduced at lengths of 36Q and 47Q (Fig. 4A). With respect to Q1 tract composition, there was general agreement in gene expression patterns (Fig. 4B) and Med15 phenotypes (Fig. 3). The non-functional RvHs and 12L Q1 inserts exhibited the most extreme changes in gene expression, followed by the partially functional 15LQ and NAB.

### The role of glutamine bias and coiled-coil propensity in Med15 Function

The extreme glutamine bias in yeast and other fungal Med15 proteins as well as in Med15 orthologs in animals [21] might suggest that there is a mechanistic basis for the overrepresentation. Mechanistically, poly-Q tracts could influence activity by affecting protein-protein interactions, either directly or indirectly; providing disorder to allow a larger set of transient interactions; providing necessary spacing between functional domains; or by providing flexibility to the protein (Fig. 6). We found that the 10 amino acid flexible spacer sequence (PGSAGSAAGG) worked as well if not better that the native 12Q sequence. These results suggest that flexibility may be more important than the sequence. The fact that residues at Q1 were not functionally constrained to be glutamine residues suggests the Q1 tract is not an interaction motif participating directly in protein-protein interactions. This is consistent with our observation that removal of the Q1 tract reduced but did not eliminate activity and that previous studies have not attributed protein interactions to the Q1 tract. In the context of the hGR reporter, two residues, Q198 and V199, downstream of the Q1 tract were found to be critical for the interaction with hGR τ fragment [18]. Gcn4, and more recently, Gal4, have been shown to interact with the ABDs of Med15 [29, 42]. While the Q1 tract is adjacent to ABD1, it is not a required component of the ABD1 interaction surface.

**Figure 6.**
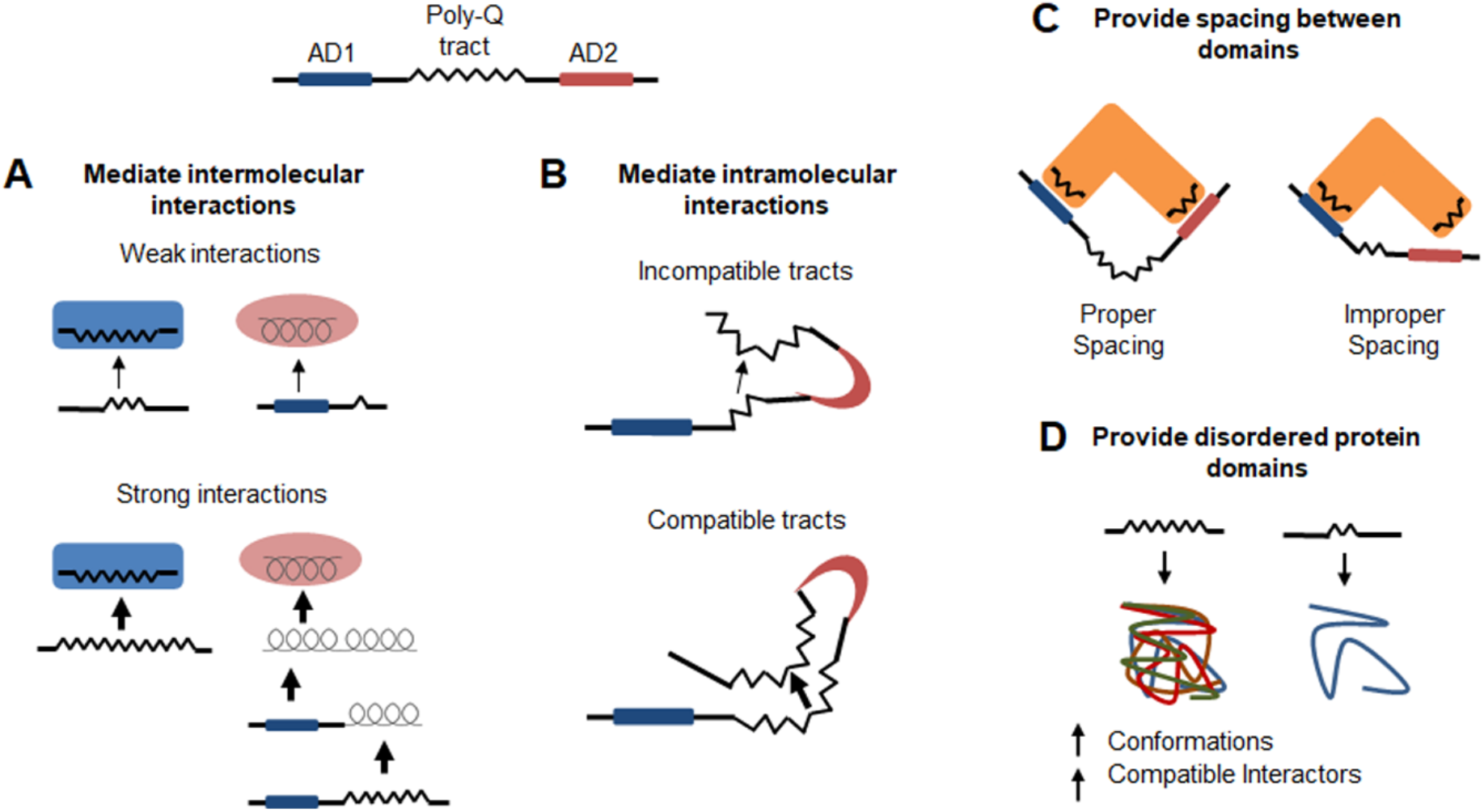
Functions of poly-Q proteins. **(A)** Poly-Q tracts can directly mediate interactions in a length-dependent manner through the formation of hydrogen bonding between helical or beta sheet structures. Poly-Q tracts can also mediate interactions indirectly by promoting coiled-coil structures in adjacent sequence which can serve as interaction domains. **(B)** The ability for poly-Q tracts to mediate interactions could influence the intramolecular dynamics of a protein with tract combinations of specific lengths being compatible. **(C)** Poly-Q tracts can function as linker domains and provide flexibility. **(D)** Poly-Q tracts promote disorder which could permit an increase in the number of structural conformations a protein can assume as well as increase the number of potential compatible interactors.

Glutamine rich sequences in Med15 and other proteins have been reported [5, 14, 16] to confer functionally important coiled-coil structure. However, we found that the presence of periodic proline residues known to perturb coiled coil structures in circular dichroism studies of Med15 related peptides [14] did not impair activity. The absence of any effect of proline residues in the Q1 tract is consistent with the naturally occurring proline interruptions in the glutamine-rich regions in functional animal orthologs of Med15 [21]. In contrast, we found that the insertion of sequences with established coiled-coil propensity such as the coiled-coil forming protein in human Med15 [16] and especially the glutamine-adjacent region from the C-terminus of the Nab3 protein [5] dampen Med15 activity (Fig. 3), suggesting that torsional flexibility of the Q1 region may be important.

Interestingly, two Q1 substitutions were not tolerated at all (Fig. 3). One was the 12L (inverted glutamine, CTG) sequence, and the other, the inverted Human Med15 coil (RvHs), which is also very leucine rich (Fig. 3). The basis for the deleterious effect of the leucine sequence is unclear, however, we previously observed that 12L insertions at the Q1 position of the KIX277-799Δ-Med15 construct were dominant negative when introduced into a wild type *MED15*^+^ strain (data not shown) suggesting that the 12L derivative may be dimerizing with native Med15 to reduce its activity.

### The role of disorder and spacing at Q1 of Med15

The possibility that the glutamine enrichment, including the poly-Q tracts, found throughout the midsection of Med15 simply promote disorder remains plausible. By promoting disorder, poly-Q tracts could permit an increase in the number of structural conformations Med15 can assume as well as increase the number of potential compatible interactors. However, our data is most consistent with a role for the Q1 tract as a spacer or hinge (Fig. 6). As seen in studies of the Huntingtin protein, poly-Q tracts may allow adjacent domains within a protein to interact [7]. Both the removal and extensive expansion of the poly-Q tract of the Huntingtin protein disrupts the proper hinge function [7]. Interestingly a shorter Huntingtin Q tract of 6 glutamines was partially defective, while a tract enriched for 12 glycine residues instead of glutamine was able to restore the hinge structure of a poly-Q deleted sequence in the Huntingtin protein [7]. The shortest Q1 tract we have identified in *MED15* from sequenced genomes is 10 glutamine residues [21]. What is not currently understood is what precise regions of the protein might interact or benefit from flexibility at the Q1 locus, but our data represent important support for environmentally dynamic conformations of Med15 within the Mediator Complex that may contribute to stress-specific transcriptomes.

## Materials and Methods

### Strains

Strains used in this study are derivatives of the S288C laboratory strain [43]. The *med15* deletion strain is from the deletion collection [44]. Phospho-site mutant strains are also S288C derived. These were a gift from Dr. Patrick Cramer and Mathias Mann [39]. Genotypes of additional strains used in this study are listed in Supplementary Table 1.

### Preparation of UAS_G_-*lacZ* Strains

*GAL4 gal80* (JF2626) and *GAL4 GAL80* (JF2624) strains with an integrated UAS_G_-*lacZ* reporter were prepared by mating the *gal4 gal80* UAS_G_-*lacZ* strain MaV103 (MATa) [45] with BY4716 (MATα) [44]. Diploids from this cross were sporulated and tetrads were analyzed to identify *GAL4 GAL80* UAS_G_-*lacZ* and *GAL4 gal80* UAS_G_-*lacZ* strains. *GAL4* strains were identified by growth on YPGal media. Strains with *lacZ* reporters were identified by production of blue color on X-Gal plates. *GAL80* strains were identified by white colony color on X-Gal plates with glucose as the carbon source and blue colony color on X-Gal plates with galactose as the carbon source.

*MED15* was deleted from both *GAL4* UAS_G_-*lacZ* strains to produce *GAL80 med15* (JF2629) and *gal80 med15* (JF2631) strains. A *med15Δ*::kanMX4 deletion cassette was amplified from genomic DNA isolated from a *med15Δ* strain (OY320) using the primers *MED15* F-245 and *MED15* R+3498. The deletion cassette was transformed into JF2624 and JF2626 using a standard lithium acetate transformation protocol with the addition of a two-hour outgrowth in YPD. Transformation reactions were plated on YPD + 350 μg/mL G418. A deletion at the *MED15* locus was verified by PCR (primers *MED15* F-245 and *MED15* R+3498) of individual transformants.

### Variant Med15 Constructs

Plasmids constructed or acquired for this study are listed in Supplementary Table 2. A series of constructs encoding synthetic Med15 proteins varying in number of domains and polyglutamine tract lengths were generated (Fig. 1). These constructs fall into four categories: internal deletions, KIX277-799Δ-Med15, KIXQ2Q3Δ-Med15, and KIXΔ-Med15 that are described below.

### *MED15* Internal Deletions

Two independent internal deletion *MED15* mutants (153-687Δ and 46-619Δ) were prepared by restriction digestion of plasmid-borne intact *MED15* alleles. Designations correspond to amino acid residues removed in the internal deletion. The 153-687Δ internal deletion plasmid (pDC2285) was prepared by digestion with *Bsu*36I and subsequent yeast homologous recombination between sequence in the Q1 and Q3 tracts. The 46-619Δ internal deletion plasmid (pDC2286) was prepared by digestion with *Eco*RI and subsequent ligation to join compatible ends.

### *MED15* Gene Fragments (gBlocks)

gBlocks (Integrated DNA Technologies) corresponding to the KIX domain (bp 1-348), Q2 region (bp 841-1650), Q3 region (bp 1651-2394), and combined Q2 Q3 region (bp 841-2394) were synthesized. In gBlocks containing the Q2 region the Q2 tract and adjacent sequence (bp 1234-1446) were removed and replaced with an *Afe*I site by incorporating the silent base change, T1233C. In gBlocks containing the Q3 region the Q3 tract and adjacent sequence (bp 2002-2115) were removed and replaced with a *Bmg*BI site by incorporating the silent base changes T2001C and G2118C. All gBlocks have 80 bp of 5’ and 3’ homology to KIX277-799Δ-*MED15* on the pRS315 M-WT plasmid. Each gBlock was A-tailed using Taq DNA polymerase (New England Biolabs), ligated into pCR2.2-TOPO (TOPO, Invitrogen) and screened on LB-Amp + X-Gal plates. Light blue and white transformants were screened by PCR using M13 F and M13 R primers. For gBlock clones that included the Q2 and/or Q3 regions internal *MED15* primers (*MED15* F+1204 and *MED15* F+1924) were used in addition to the M13 primers to reduce the size of the amplified fragment. Finally, the plasmids were confirmed by sequencing and each clone was stored (GC1012, GC1013, GC1014, and GC1025).

A Q1 region (Q1R; bp 346-831) fragment was constructed with the Q1 tract replaced by an *Afe*I site using two step PCR. First, sequence on either side of the Q1 tract was amplified with overlapping forward and reverse primers having an *Afe*I site between Q1 flanking sequence (0Q F and 0Q R) paired with reverse (Q1R gap repair R) and forward (Q1R gap repair F) primers respectively. Next, the two PCR products and the outer forward and reverse primers were used to amplify a 0-Q1:*Afe*I Q1R fragment. This construct was ligated into the TOPO vector for storage and propagation (pDC2178).

gBlock DNA was digested out of the TOPO vectors using *Bst*XI (New England Biolabs), sites which are present in the vector on either side of the insert and are absent from gBlocks. The gBlock band was extracted and purified from agarose gels using the QIAquick Gel Extraction Kit (Qiagen). Alternatively, the KIX gBlock was PCR amplified from the TOPO vector using KIX gBlock F and KIX gBlock R primers.

### KIXQ2Q3Δ-Med15 Variants

KIXQ2Q3Δ-*MED15* constructs are intermediate in size relative to the KIX277-799Δ-*MED15* and full length *MED15* in that the KIX domain is still absent as it is in the KIX277-799Δ-Med15 while the central Q-rich region of Med15 is present (Fig. 1). These constructs were made by the addition of the Q2Q3 gBlock to either the 0-Q1 or 12-Q1 KIX277-799Δ-Med15 construct. The Q1R region of pRS315 M-WT, 0-Q1 KIX277-799Δ-*MED15*, and 12-Q1 KIX277-799Δ-*MED15* were PCR amplified using Q1R gap repair F and Q1R gap repair R. These PCR products were individually co-transformed with the Q2Q3 gBlock and *Pst*I digested pRS315 M-WT. The DNA fragment pairs have sequence overlap allowing for homologous recombination with each other as well as with the gapped plasmid. Two versions of 12-Q1 KIXQ2Q3Δ-*MED15* were constructed by using either pRS315 M-WT Q1R PCR (pDC2149) or using 12-Q1 KIX277-799Δ-*MED15* Q1R PCR (pDC2138). 0-Q1 KIXQ2Q3Δ-*MED15* (pDC2136) was constructed by using 0-Q1 KIX277-799Δ-*MED15* Q1R PCR.

Variant Q1 KIXQ2Q3Δ-*MED15* constructs were prepared by first introducing natural or synthetic Q1 sequences into the *Afe*I site in pDC2178 and then gap-repairing partially *Afe*I digested pDC2136 with a Q1R PCR product amplified from the pDC2178 derivative using the primers Q1R gap repair F and Q1R gap repair R. Natural Q1 tract length variants (19-Q1, pDC2150; and 21-Q1, pDC2151) were constructed by amplifying the Q1R sequence from pooled genomic DNA from multiple wine yeast strains using Q1R gap repair F and Q1R gap repair R primers. Synthetic Q1 tract length variants (12-Q1, pDC2149; 24-Q1, pDC2144; 36-Q1, pDC2146) were constructed by ligation of glutamine tract coding duplex DNA (5’-CAACAACAACAACAACAACAGCAGCAGCAGCAACAG). This method produced tracts of glutamine codons in multiples of 12. This method also produced tracts of polyleucine when the duplex was ligated in the reverse orientation (12L-Q1, pDC2293). A synthetic variant (47-Q1, pDC2147) was constructed by mismatch recombination between two tracts of 36 glutamine codons. A short Q tract (6-Q1, pDC2260), a non-Q spacer (Spacer-Q1, pDC2185), modified Q tracts (12PQ-Q1, pDC2262; 24PQ-Q1 pDC2263; 15LQ-Q1, pDC2291), and heterologous sequences (NAB3-Q1, pDC2292; FrHs-Q1, pDC2294; RvHs-Q1, pDC2295) were constructed by ligation of duplexed oligos. Duplexed oligos used in the ligation included: a glycine rich spacer sequence (5’-CCAGGTTCTGCTGGTTCTGCTGCTGGTGGT: PGSAGSAAGG); a short Q tract of 6Q (5’-CAACAACAACAACAGCAG); a coiled-coil disrupting sequence (5’-CCTCAACAACAGCCTCAGCAACCACAGCAACAACCA: PQQQPQQPQQQP); a coiled-coil promoting sequence (5’-TTGCAACAACAGTTACAGCAATTGCAGCAACAACTGTTGTTGCAA: LQQQLQQLQQQLLLQ); the C-terminal helix of Nab3 (5’-AATGTTCAAAGTCTATTAGATAGTTTAGCAAAACTACAAAAG: NVQSLLDSLAKLQK); and a portion of the human Med15 Q tract (5’-CTGCAGCTCCAGCAGGTGGCGCTGCAGCAGCAGCAGCAACAGCAGCAGTTCCAGCA GCAG) ligated in the forward (LQLQQVALQQQQQQQQFQQQ) and reverse orientation (LLLELLLLLLLLQRHLLELQ).

### KIXΔ-Med15

Plasmid KIX277-799Δ-*MED15* (pRS315 M-WT) was digested with *Bst*API to create a gap between the Q1 containing region (Q1R) and the MAD region. The gap was repaired using the Q2/Q3 gBlock isolated from GC1014 to generate pDC2149 (KIXQ2Q3Δ-Med15). pDC2149 was sequentially digested at the Q3 locus using *Bmg*BI and gap repaired using the Q3 PCR fragment (primers *MED15* F+1682 and *MED15* R+2385) from the lab strain BY4742 and then digested at the Q2 locus using *AfeI* and gap repaired using the Q2 PCR fragment (primers *MED15* F+854 and *MED15* R+1781) from the lab strain BY4742 to generate pDC2217 (KIXΔ-Med15).

### Split Ubiquitin

The first 813 bp of the *MSN2* ORF were fused upstream of the C-ubiquitin sequence of the P_*MET17*_ C-ubiquitin R*URA3* plasmid (addgene 131163, [46]) to create pDC2279. A P_*CUP1*_ N-ubiquitin plasmid was constructed by amplifying the *CUP1* promoter through N-ubiquitin sequence from the integration plasmid (addgene 131169, [46]) and subsequently the terminator sequence from *MED15* was added downstream to create pDC2278. Fragments of the *MED15* gene were fused upstream of the N-ubiquitin sequence (KIX, pDC2283; Q1R, pDC2284; KIX+Q1R, pDC2282; Q1R-NAB, pDC2280; Q1R-Hs-rev, pDC2281; Q1R-0Q, pDC2301; Q1R-24Q, pDC2302; Q1R-Spacer, pDC2303; Q1R-6Q, pDC2304; Q1R-12PQ, pDC2305; Q1R-24PQ, pDC2307; Q1R-12L, pDC2307).

### Yeast Methods

#### Colony PCR

Yeast colonies were used for rapid PCR screening using Taq polymerase, colony PCR buffer (final concentrations: 12.5 mM Tris-Cl (pH 8.5), 56 mM KCl, 1.5 mM MgCl_2_, 0.2 mM dNTPs), and primers at a final concentration of 0.2 μM. Reaction mixes were aliquoted into PCR tubes and then a small amount of yeast cells were transferred from a streak plate into each tube using the end of a 200 μL micropipet tip. A standard hot-start thermocycler program with extension at 68°C was adjusted for the Tm of the primer set and size of the amplification product.

#### Yeast Transformation

Transformations of *med15Δ* strains were conducted using the Frozen-EZ Yeast Transformation II kit (Zymo Research) with modifications. 10 mL of early log phase cells were pelleted and washed with 2 mL EZ 1 solution, repelleted and resuspended in 1 mL EZ 2 solution. Aliquots were frozen at -80°C. 0.2-1 μg DNA and 100 μL EZ 3 solution were added to 10 μL competent cells for each transformation. Transformations were plated on selective media following incubation at 30°C for 45 minutes. Transformations into all other strains were conducted using a standard lithium acetate transformation protocol [47, 48].

### Media and Phenotype Testing

Cultures were grown in rich media (YPD) or synthetic complete media lacking single amino acids (SC-Leu or SC-Ura). Spot assays were performed on rich media or synthetic complete media with additives.

Exponentially growing subcultures were diluted differentially to achieve a consistent initial concentration of 5×10^6^ cells/mL. 5-fold or 10-fold serial dilutions were carried out in the wells of a sterile 96-well microtiter dish. 2 μL volumes of each dilution were spotted on different types of solid media. Plates were incubated at desired temperatures (22°C, 30°C, and 38°C). Phenotypes were observed and imaged daily using a flatbed scanner.

Spot assays were quantified using ImageJ. Grayscale TIFF formatted images of plates were imported into ImageJ. Background pixels, that did not correspond to yeast colonies, were set to an intensity of 0 using an intensity threshold cutoff. Threshold cutoffs determined for each plate separately. An area containing yeast colonies was selected using the rectangle tool. The same rectangular area selection was used across a single experiment. For each strain the percent area was measured (pixels corresponding to colonies/total area * 100). Quantification for each strain on each condition normalized to the growth of that strain on YPD.

A microtiter dish based chemical sensitivity assay was implemented with minor modifications to the method previously described [49] to achieve increased resolution. Cells of each genotype in biological triplicate (transformants) were grown to saturation in SC-Leu media, diluted to an OD_600_ of 0.1-0.2 and inoculated 1:1 into 2xYPD with various concentrations of drug or in alternative types of media and allowed to incubate overnight at the indicated temperature. OD_600_ readings were conducted using a Cytation microplate reader following 30 seconds of shaking at an endpoint of 20-24 hours. Relative growth was calculated after background subtraction by dividing the absorbance for each treated well with the corresponding value of the 0-drug control well. Relative growth for each genotype was plotted for a single drug concentration chosen for causing a reduction in growth of the wild type *MED15* strain of less than 50%.

β-Galactosidase Reporter Assays

### Human Glucocorticoid Receptor Tau 1 Fragment Dependent Reporter

Yeast carrying an expression vector for the human glucocorticoid receptor Tau 1 fragment transcription factor (hGR) and the glucocorticoid receptor driven *lacZ* reporter (JF2768) were transformed with a third plasmid carrying the desired *MED15* allele (pRS315 M-WT derivative) or an empty vector without a *MED15* allele (pRS315). As previously described, transformants were prescreened on X-Gal plates to identify transformants displaying the “average” amount of reporter activity for that strain [18]. Outliers that were either noticeably more or less blue than the other transformants were excluded.

At least 3 transformants per strain were grown in minimal media (SD+Ade+Lys) to a concentration of approximately 2×10^7^ cells/mL. To induce the hGR transcription factor, CuSO_4_ was added to log phase cultures to a final concentration of 0.25 mM and the cultures were grown for an additional hour. Cultures were then prepared for protein extraction by pelleting the yeast, adding glass beads and extraction buffer, and freezing at -80°C. Analysis was conducted over the next couple days to avoid degradation or deterioration of the sample.

### Gal4 Dependent Reporter

*med15Δ gal80Δ* strain (JF2631) containing an integrated UAS_G_-*lacZ* reporter was transformed with plasmids containing the desired *MED15* allele or an empty vector. Transformants were subcultured in SC-Leu 2% raffinose and grown at 30°C to log phase (1-2×10^7^ cells/mL). *med15Δ GAL80* strains (JF2629) containing an integrated UAS_G_-*lacZ* reporter were likewise cultured in SC-Leu 2% raffinose to saturation but then subcultured in SC-Leu with 2% raffinose + 2% glucose or SC-Leu 2% raffinose + 2% galactose and grown at 30°C to log phase (1-2×10^7^ cells/mL). Cultures prepared for protein extraction as above and stored at -80°C.

### Protein Extraction

Yeast cells were harvested from log-phase cultures in reporter-specific growth conditions and frozen at -80°C. Pelleted yeast cells were resuspended in Breaking Buffer (0.1 M Tris pH 8, 20% glycerol, 1 mM DTT) in tubes containing glass beads (200 mg, 0.4 mm) and were lysed using a TissueLyser LT (Qiagen). To minimize protein degradation, 5 μL of 40 mM PMSF was added periodically throughout the process of preparing extracts. Tubes were alternately shaken at 50 oscillations/second for 20-30 seconds and then placed on ice for at least 1 minute for a total of 2 minutes of shaking. Cell debris was pelleted at 13,000 rpm for 10 minutes at 4°C. The supernatant was transferred to a new tube containing additional PMSF. Protein extracts were stored at -80°C or kept on ice until used for subsequent analyses.

### Protein Concentration Determination (Bradford Assay)

Total protein concentrations were determined using a modified Bradford Assay (BioRad). 2-10 μL of each protein extract or BSA standard (100-1000 mg/mL) was added into at least two wells of a 96-well plate. Several wells were left empty to serve as blanks. To each well, 200 μL of Bradford assay working solution was added using an 8-tip multichannel pipet. A standard curve was generated using a series of BSA standards of known concentrations designed to produce absorbance readings within the linear range of the spectrophotometer. Final absorbance readings were taken after 15 minutes at a wavelength of 595 nm. The average absorbance of the blank wells was subtracted from all readings. A best fit linear regression based on the standard calibration curve was used to determine sample concentration.

### Enzymatic Assay

Approximately 10 ng of total protein (5-50 μL protein extract) was diluted to 1 mL in Z-buffer (60 mM Na_2_HPO_4_, 40 mM NaH_2_PO_4_, 10 mM KCl, 1 mM MgSO_4_, 50 mM β-mercaptoethanol (BME)) in 13 mm glass tubes and incubated in a temperature block held at 28°C. 200 μL ONPG (4 mg/mL in Z-buffer; made fresh for each experiment) was added to each tube (time 0). Reactions were stopped once the solutions turned yellow by addition of 500 μL of 1 M Na_2_CO_3_. A_420_ measurements were taken for each sample.

### Data Processing

Beta-galactosidase values were calculated using the formula:

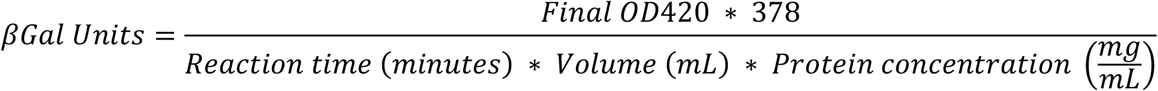

Due to systematic difference between experiments, such as different ONPG solution preparations or quality of extracts, relative expression values were calculated for each experiment to be able to compare strains measured in different experiments. When necessary, the average β-gal value for the wild type strain was set to 100% for each experiment.

### Split Ubiquitin Assay

Yeast strains containing both an N-ubiquitin and a C-ubiquitin-R*URA3* expressing plasmids were cultured in SD+LysUra to log-phase at 30°C to a concentration of ∼1×10^7^ cells/mL. CuSO_4_ was added to a final concentration of 0.25mM and the cultures were incubated for an additional hour. Cultures were serially diluted and 2 μL was plated on media without uracil and media containing 5-FOA.

### RNA Methods

Yeast strains were cultured in SC-Leu to saturation and then subcultured in 10 mL YPD and grown to log-phase at 30°C to a concentration of ∼2×10^7^ cells/mL. After incubation yeast were pelleted, washed with cold water, frozen with dry ice, and stored at -80°C. RNA was extracted using a hot acid phenol protocol with modifications [50]. Pellets were resuspended in 400 μL TES solution (10 mM Tris, 10 mM EDTA, 0.5% SDS) and an equal volume of acid phenol was added to each tube and vortexed for 10 seconds. Tubes were incubated in a 65°C water bath for 1 hour and vortexed every 10-12 minutes. Phases were separated by microcentrifugation in the cold for 5 minutes. The aqueous layer was extracted into a new tube avoiding the DNA-enriched interface. A second round of acid phenol extraction was conducted followed by a chloroform extraction. RNA was precipitated with 300 μL of 4 M LiCl in dry ice for 20 minutes followed by 5 minutes of high-speed centrifugation at 4°C. The pellet was washed with 500 μL ice cold 70% ethanol and then dried for 10 minutes at 37°C. RNA pellets were resuspended in DEPC treated water and stored at -20°C.

Contaminating genomic DNA was removed using the DNase Max Kit (Qiagen). 10 μL of a 1x Master Mix (5 μL 10X Buffer, 5 μL water, and 0.5 μL DNase I) was added to 40 μL RNA. Tubes were incubated at 37°C for 30 minutes. DNase was removed by addition of 5 μL of the DNase removal resin and room temperature incubation for 10 minutes with periodic agitation. The resin was pelleted by centrifugation for one minute at the highest setting of a microcentrifuge. The supernatant containing DNA-free RNA was transferred to a new tube. The concentration of the purified RNA was determined using a NanoDrop spectrophotometer (Thermo Scientific).

### cDNA First Strand Synthesis

cDNA was prepared using Superscript III First-Strand Synthesis System (Invitrogen) or Lunascript (New England Biolabs). Transcripts were amplified with either random hexamer (Invitrogen) or anchored oligo-dT20 (Integrated DNA Technologies) primers. For anchored oligo-dT20 primers 1 μg RNA, 50 ng primer, and 0.01 μmol dNTPs were mixed in a final volume of 10 μL. Tubes were incubated for 5 minutes at 65°C and then 2 minutes on ice. 10 μL cDNA Master Mix (2x RT buffer, 10 mM MgCl_2_, 0.02 M DTT, 40 U RNase OUT, 200 U Superscript III Reverse Transcriptase) or mock Master Mix which did not contain reverse transcriptase was added to each tube which were then incubated for 60 minutes at 50°C and 5 minutes at 85°C. To degrade RNA, 2 U RNase H was added to each tube and then the tubes were incubated for 20 minutes at 37°C. For random hexamer primers tubes were incubated 10 minutes at 25°C and then 50 minutes at 50°C.

### qRT-PCR

Transcript abundance was quantified for each sample relative to a normalization transcript using the PerfeCTa SYBR Green FastMix (Quantabio) in 96-well PCR plates (Hard-Shell 480 PCR Plates, Bio-Rad) measured in a LightCycler 480 (Roche). Individual 10 μL reactions consisted of 5 μL FastMix, 1 μL target-specific primer pairs (0.5 μL 10 μM forward primer and 0.5 μL 10 μM reverse primer), 3 μL DEPC treated water, and 1 μL cDNA, mock cDNA, or water. Plates were sealed with optically clear film (PlateSeal). A standard SYBR green PCR program was used with modifications: 95°C for 5 minutes, 45 cycles of 95°C for 10 seconds, 55°C for 10 seconds, and 72°C for 20 seconds with a single fluorescence acquisition. A melting curve was conducted directly following the PCR program to confirm the presence of individual species amplified in each well. The melting curve program was 95°C for 5 seconds, 65°C for 1 minute, and ramp up to 97°C by 0.11°C/s with continuous fluorescence acquisition. *ALG9* was used as a normalization transcript. All samples were measured with three technical replicates. Mock cDNA (prepared without the addition of reverse transcriptase) and water were used as negative controls.

### Data Analysis

The relative abundance of target transcripts was determined by calculating the crossing point (CP) or cycle threshold (CT) value for each reaction. CP values were calculated using the Second Derivative Maximum method implemented in the LightCycler 480 Software (Roche). CT values were calculated using the Comparative CT (ΔΔCT) method implemented in the QuantStudio 3 Software (Applied Biosystems). The average CP or CT for technical replicates amplifying the target transcript with a specific RNA sample was normalized to the average CP or CT for technical replicates amplifying the normalization transcript. This ratio was then compared across samples as a depiction of relative abundance of the target transcript in each sample.

### Primers

Primers used in this study are listed in Supplementary Table 3. All primers were synthesized by Integrated DNA Technologies. The use of each primer in construction, screening, and sequencing is noted in Supplementary Table 3. *MED15* alleles were submitted to the Roy J. Carver Center for Genomics (CCG) for Sanger sequencing to confirm the gene sequence and identify SNPs. All primer pairs used to amplify transcripts in qRT-PCR are listed in Supplementary Table 4. Primer pair efficiency was measured by qRT-PCR analysis of serial dilution of control RNA.

## Acknowledgements

The authors gratefully acknowledge the generous gift of strains and plasmids from Dr. Patrick Cramer and Mathias Mann, (phospho-site mutant strains), Dr. Young Chul Lee (glucocorticoid reporter plasmids), and Nils Johnsson via addgene (split ubiquitin vectors). The authors also appreciate comments and suggestions from Dr. Daniel Weeks and Dr. Bryan Phillips and from members of the lab.

## Funding

Funding for this project was from T32 (Bioinformatics Training Grant), a University of Iowa CLAS Dissertation Writing Fellowship, and the University of Iowa, Department of Biology Evelyn Hart Watson Summer Fellowship, and supplemental funding to NIH award R35 GM058939-19S1.

## Notes

### Competing Interest Statement

The authors have declared no competing interest.

## References

1. Paulson, H.L. and K.H. Fischbeck, Trinucleotide repeats in neurogenetic disorders. Annu Rev Neurosci, 1996. 19: p. 79–107.

2. Gatchel, J.R. and H.Y. Zoghbi, Diseases of unstable repeat expansion: mechanisms and common principles. Nat Rev Genet, 2005. 6(10): p. 743–55.

3. Hands, S., C. Sinadinos, and A. Wyttenbach, Polyglutamine gene function and dysfunction in the ageing brain. Biochim Biophys Acta, 2008. 1779(8): p. 507–21.

4. Stott, K., et al., Incorporation of glutamine repeats makes protein oligomerize: implications for neurodegenerative diseases. Proc Natl Acad Sci U S A, 1995. 92(14): p. 6509–13.

5. Loya, T.J., T.W. O’Rourke, and D. Reines, Yeast Nab3 protein contains a self-assembly domain found in human heterogeneous nuclear ribonucleoprotein-C (hnRNP-C) that is necessary for transcription termination. J Biol Chem, 2013. 288(4): p. 2111–7.

6. Garbett, K.A., et al., Yeast TFIID serves as a coactivator for Rap1p by direct protein-protein interaction. Mol Cell Biol, 2007. 27(1): p. 297–311.

7. Caron, N.S., et al., Polyglutamine domain flexibility mediates the proximity between flanking sequences in huntingtin. Proc Natl Acad Sci U S A, 2013. 110(36): p. 14610–5.

8. Nucifora, F.C., Jr., et al., Interference by huntingtin and atrophin-1 with cbp-mediated transcription leading to cellular toxicity. Science, 2001. 291(5512): p. 2423–8.

9. Schaefer, M.H., E.E. Wanker, and M.A. Andrade-Navarro, Evolution and function of CAG/polyglutamine repeats in protein-protein interaction networks. Nucleic Acids Research, 2012. 40(10): p. 4273–4287.

10. Caprioli, M., et al., Clock gene variation is associated with breeding phenology and maybe under directional selection in the migratory barn swallow. PLoS One, 2012. 7(4): p. e35140.

11. Bryan, A.C., et al., A Variable Polyglutamine Repeat Affects Subcellular Localization and Regulatory Activity of a Populus ANGUSTIFOLIA Protein. G3 (Bethesda), 2018. 8(8): p. 2631–2641.

12. Rice, C., et al., The Nature, Extent, and Consequences of Genetic Variation in the opa Repeats of Notch in Drosophila. G3 Genes|Genomes|Genetics, 2015. 5(11): p. 2405–2419.

13. Gemayel, R., et al., Variable Glutamine-Rich Repeats Modulate Transcription Factor Activity. Mol Cell, 2015. 59(4): p. 615–27.

14. Fiumara, F., et al., Essential role of coiled coils for aggregation and activity of Q/N-rich prions and PolyQ proteins. Cell, 2010. 143(7): p. 1121–35.

15. Loya, T.J., et al., A network of interdependent molecular interactions describes a higher order Nrd1-Nab3 complex involved in yeast transcription termination. J Biol Chem, 2013. 288(47): p. 34158–67.

16. Batlle, C., et al., MED15 prion-like domain forms a coiled-coil responsible for its amyloid conversion and propagation. Commun Biol, 2021. 4(1): p. 414.

17. Barberis, A., et al., Contact with a component of the polymerase II holoenzyme suffices for gene activation. Cell, 1995. 81(3): p. 359–68.

18. Kim, D.H., et al., Functional conservation of the glutamine-rich domains of yeast Gal11 and human SRC-1 in the transactivation of glucocorticoid receptor Tau 1 in Saccharomyces cerevisiae. Mol Cell Biol, 2008. 28(3): p. 913–25.

19. Nishizawa, M., S. Taga, and A. Matsubara, Positive and negative transcriptional regulation by the yeast GAL11 protein depends on the structure of the promoter and a combination of cis elements. Mol Gen Genet, 1994. 245(3): p. 301–12.

20. Herbig, E., et al., Mechanism of Mediator recruitment by tandem Gcn4 activation domains and three Gal11 activator-binding domains. Mol Cell Biol, 2010. 30(10): p. 2376–90.

21. Cooper, D.G. and J.S. Fassler, Med15: Glutamine-Rich Mediator Subunit with Potential for Plasticity. Trends Biochem Sci, 2019. 44(9): p. 737–751.

22. Thakur, J.K., et al., A nuclear receptor-like pathway regulating multidrug resistance in fungi. Nature, 2008. 452(7187): p. 604–9.

23. Thakur, J.K., et al., Mediator subunit Gal11p/MED15 is required for fatty acid-dependent gene activation by yeast transcription factor Oaf1p. J Biol Chem, 2009. 284(7): p. 4422–8.

24. Nogi, Y. and T. Fukasawa, A novel mutation that affects utilization of galactose in Saccharomyces cerevisiae. Curr Genet, 1980. 2(2): p. 115–20.

25. Traven, A., B. Jelicic, and M. Sopta, Yeast Gal4: a transcriptional paradigm revisited. EMBO Rep, 2006. 7(5): p. 496–9.

26. Hinnebusch, A.G. and G.R. Fink, Positive regulation in the general amino acid control of Saccharomyces cerevisiae. Proc Natl Acad Sci U S A, 1983. 80(17): p. 5374–8.

27. Natarajan, K., et al., Transcriptional profiling shows that Gcn4p is a master regulator of gene expression during amino acid starvation in yeast. Mol Cell Biol, 2001. 21(13): p. 4347–68.

28. Brzovic, P.S., et al., The acidic transcription activator Gcn4 binds the mediator subunit Gal11/Med15 using a simple protein interface forming a fuzzy complex. Mol Cell, 2011. 44(6): p. 942–53.

29. Tuttle, L.M., et al., Mediator subunit Med15 dictates the conserved “fuzzy” binding mechanism of yeast transcription activators Gal4 and Gcn4. Nat Commun, 2021. 12(1): p. 2220.

30. Boija, A., et al., Transcription Factors Activate Genes through the Phase-Separation Capacity of Their Activation Domains. Cell, 2018. 175(7): p. 1842–1855 e16.

31. Lallet, S., et al., Role of Gal11, a component of the RNA polymerase II mediator in stress-induced hyperphosphorylation of Msn2 in Saccharomyces cerevisiae. Mol Microbiol, 2006. 62(2): p. 438–52.

32. Gallagher, J.E.G., et al., The Polymorphic PolyQ Tail Protein of the Mediator Complex, Med15, Regulates the Variable Response to Diverse Stresses. Int J Mol Sci, 2020. 21(5).

33. Cooper, D.G., et al., Possible Role for Allelic Variation in Yeast MED15 in Ecological Adaptation. Front Microbiol, 2021. 12: p. 741572.

34. Park, J.M., et al., In vivo requirement of activator-specific binding targets of mediator. Mol Cell Biol, 2000. 20(23): p. 8709–19.

35. Thakur, J.K., et al., Mediator subunit Gal11p/MED15 is required for fatty acid-dependent gene activation by yeast transcription factor Oaf1p. The Journal of biological chemistry, 2009. 284(7): p. 4422–8.

36. Jedidi, I., et al., Activator Gcn4 employs multiple segments of Med15/Gal11, including the KIX domain, to recruit mediator to target genes in vivo. J Biol Chem, 2010. 285(4): p. 2438–55.

37. Hidalgo, P., et al., Recruitment of the transcriptional machinery through GAL11P: structure and interactions of the GAL4 dimerization domain. Genes Dev, 2001. 15(8): p. 1007–20.

38. Qi, Y., et al., Mediator Engineering of Saccharomyces cerevisiae To Improve Multidimensional Stress Tolerance. Appl Environ Microbiol, 2022. 88(8): p. e0162721.

39. Miller, C., et al., Mediator phosphorylation prevents stress response transcription during non-stress conditions. J Biol Chem, 2012. 287(53): p. 44017–26.

40. Wittke, S., et al., Probing the molecular environment of membrane proteins in vivo. Mol Biol Cell, 1999. 10(8): p. 2519–30.

41. Sadeh, A., et al., Conserved motifs in the Msn2-activating domain are important for Msn2-mediated yeast stress response. J Cell Sci, 2012. 125(Pt 14): p. 3333–42.

42. Tuttle, L.M., et al., Gcn4-Mediator Specificity Is Mediated by a Large and Dynamic Fuzzy Protein-Protein Complex. Cell Rep, 2018. 22(12): p. 3251–3264.

43. Mortimer, R.K. and J.R. Johnston, Genealogy of principal strains of the yeast genetic stock center. Genetics, 1986. 113(1): p. 35–43.

44. Brachmann, C.B., et al., Designer deletion strains derived from Saccharomyces cerevisiae S288C: a useful set of strains and plasmids for PCR-mediated gene disruption and other applications. Yeast, 1998. 14(2): p. 115–32.

45. Vidal, M., et al., Reverse two-hybrid and one-hybrid systems to detect dissociation of protein-protein and DNA-protein interactions. Proc Natl Acad Sci U S A, 1996. 93(19): p. 10315–20.

46. Dunkler, A., J. Muller, and N. Johnsson, Detecting protein-protein interactions with the Split-Ubiquitin sensor. Methods Mol Biol, 2012. 786: p. 115–30.

47. Ito, H., et al., Transformation of intact yeast cells treated with alkali cations. J Bacteriol, 1983. 153(1): p. 163–8.

48. Gietz, R.D. and R.A. Woods, Transformation of yeast by lithium acetate/single-stranded carrier DNA/polyethylene glycol method. Methods Enzymol, 2002. 350: p. 87–96.

49. Hung, C.W., et al., A simple and inexpensive quantitative technique for determining chemical sensitivity in Saccharomyces cerevisiae. Sci Rep, 2018. 8(1): p. 11919.

50. Collart, M.A. and S. Oliviero, Preparation of yeast RNA. Curr Protoc Mol Biol, 2001. Chapter 13: p. Unit13 12.

