## Supplemental Figures and Tables for "The Role of Polyglutamine in Inter- and Intra-molecular Interactions in Med15-dependent Regulation"

<u>SUPPLEMENTARY FILE CONTENTS</u>

1. SUPPLEMENTARY FIGURES 1-3
2. SUPPLEMENTARY TABLES
3. SUPPLEMENTARY REFERENCES

A

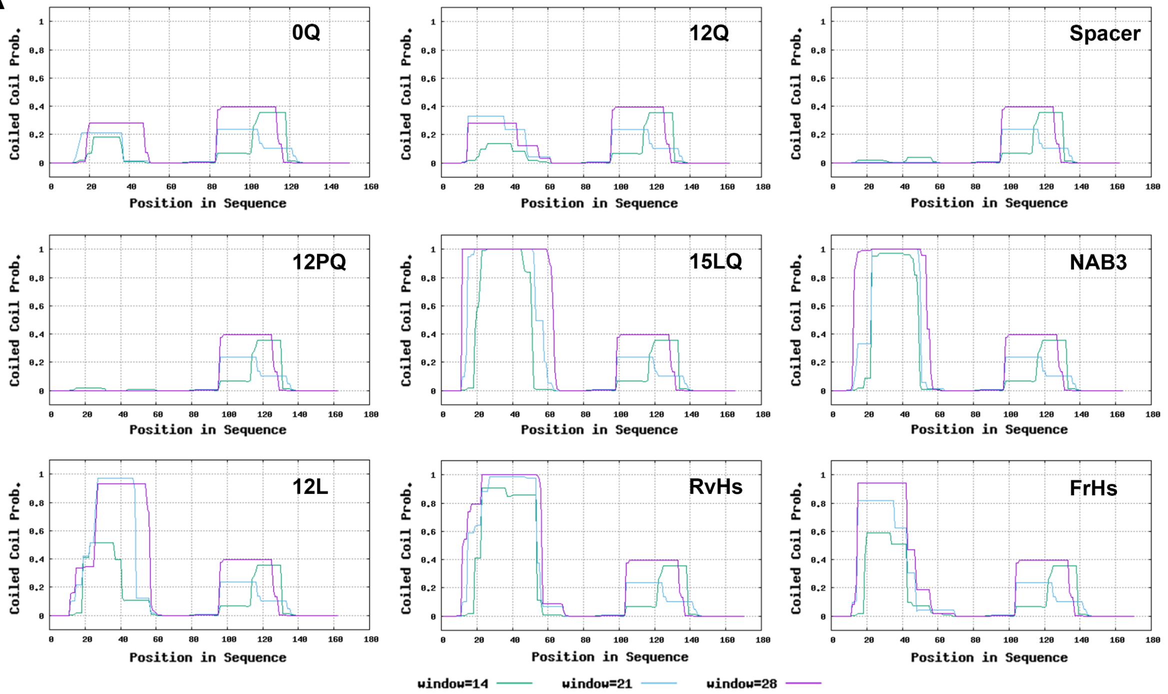


B

| Construct | Max Prob. | Area Under Curve | Residues in Q1 Peak | Average Prob. in Q1 Peak | Relative Average |
|---|---|---|---|---|---|
| **0Q** | 0.28 | 8.23 | 37.00 | 0.22 | 1.04 |
| **12Q** | 0.28 | 9.63 | 45.00 | 0.21 | 1.00 |
| **Spacer** | 0.00 | 0.00 | 0.00 | 0.00 | 0.00 |
| **12PQ** | 0.00 | 0.00 | 0.00 | 0.00 | 0.00 |
| **15LQ** | 1.00 | 51.00 | 57.00 | 0.89 | 4.18 |
| **NAB3** | 1.00 | 41.59 | 55.00 | 0.76 | 3.53 |
| **12L** | 0.93 | 32.60 | 58.00 | 0.56 | 2.63 |
| **RvHs** | 1.00 | 42.74 | 66.00 | 0.65 | 3.03 |
| **FrHs** | 0.94 | 30.99 | 59.00 | 0.53 | 2.45 |

**Supplementary Figure 1. Predicted coiled-coil structure of the Q1 locus. (A)** Coiled-coil structure of Med15 Q1R (amnio acids 116-277) with different Q1 sequences predicted using PCOILS [1]. The Q1 locus in Med15 (amino acids 147-158) corresponds to residues 32-43 of the input sequence for the 12Q construct. **(B)** Quantification of Coiled-coil predictions for the Q1 locus. Quantification of the max probability (Max Prob.), sum of all probability values for the peak (Area Under Curve), and number of residues in the Q1 peak are based on the 28-residue window for the Q1 region. The Average Probability in the Q1 Peak was calculated as the area under the curve divided by the number of residues in the Q1 peak. The average values were then made relative to the 12Q average probability (Relative Average).

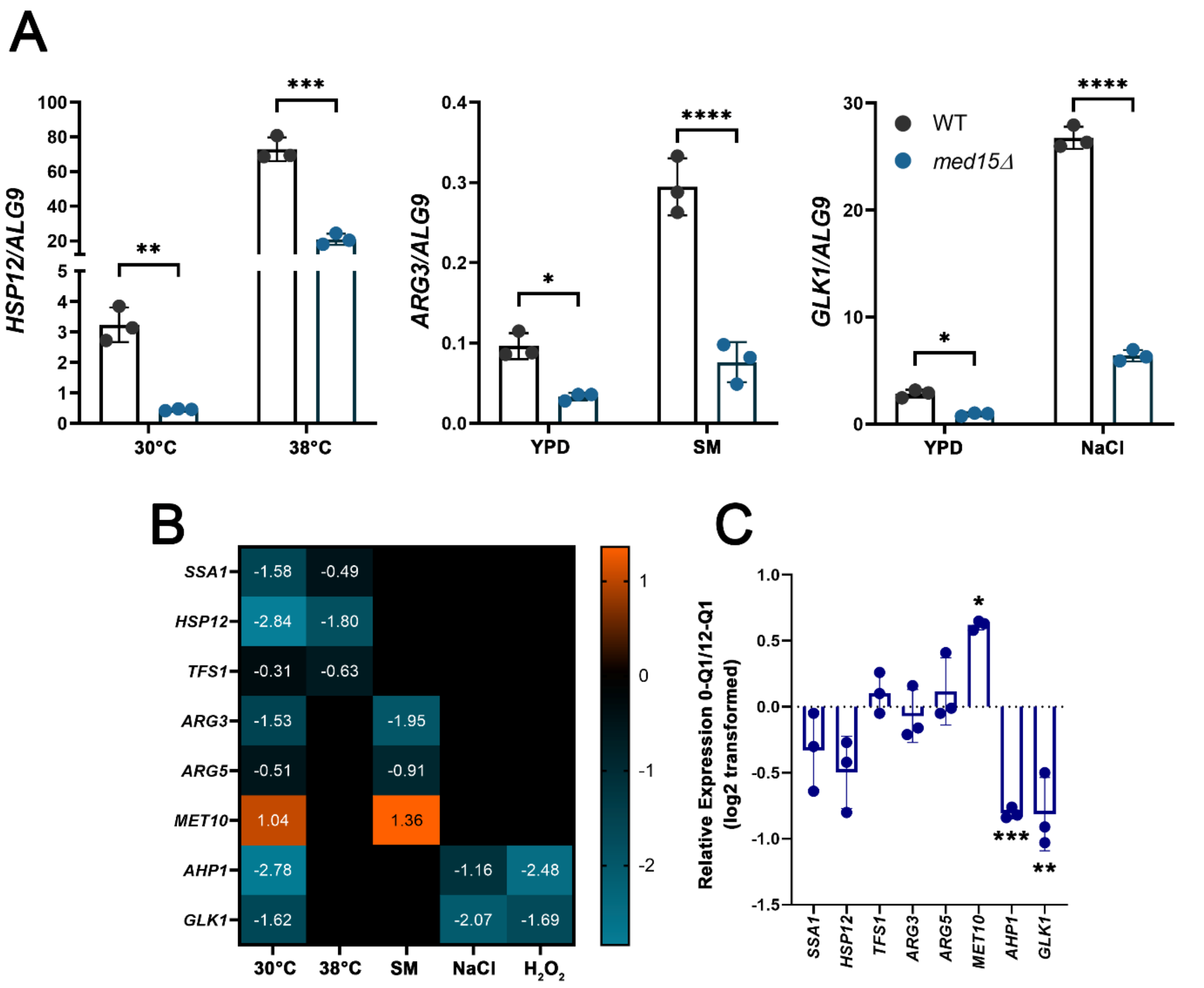


**Supplementary Figure 2. Med15-dependent basal and stress-induced expression of Gcn4 and Msn2 regulated genes.** Yeast grown to log phase in YPD at 30°C and then either maintained in that condition (30°C) or transferred to YPD: prewarmed to 38°C (38°C), containing 1 µg/mL sulfometuron-methyl (SM), containing 0.7 M NaCl (NaCl), or containing 0.5 mM $H_2O_2$ ($H_2O_2$). For each strain the target gene expression was normalized to *ALG9* and averaged over 3 biological replicates (transformants). **(A)** Expression of target genes *HSP12*, *ARG3*, and *GLK1* normalized to *ALG9* for log-phase cultures of the *med15Δ* strain (OY320) carrying a CEN plasmid with *MED15* (black) or without (blue) in YPD grown at 30°C or following treatment with heat (38°C), SM, or NaCl. **(B)** Log2 transformed ratio of target gene expression in *med15Δ* relative to WT. The ratio of gene expression was determined by dividing the average expression in *med15Δ* to the average expression in WT. **(C)** Log2 transformed relative expression of target genes in 0-Q1 vs. 12-Q1 KIXQ2Q3Δ-*MED15* 30°C YPD cultures.

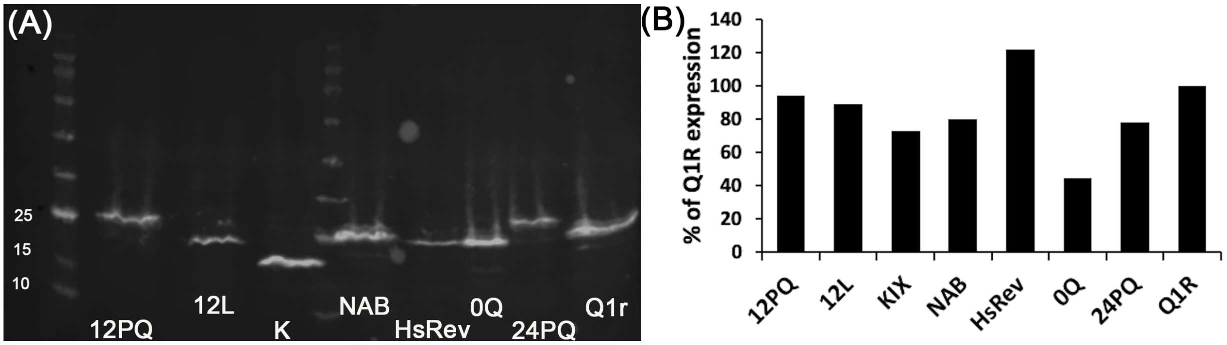


**Supplementary Figure 3. Protein production from Med15 expressing split-ubiquitin constructs.** HA Western of N-Ubiquitin-Med15 split ubiquitin constructs. **(A)** HA signal for extracts prepared using the glass bead method from 10 mL cultures grown in SC-Leu and induced for 90 min with 0.1 mM $CuSO_4$. Approximately 30 μg of protein were loaded per lane of an SDS 4-12% Tris-glycine polyacrylamide gel and transferred to an activated PVDF membrane. The membrane was blocked for 20 minutes with StartingBlock (Thermofisher) and incubated overnight at $4^{o}$ C with anti-HA antibody (1:3000 in TBST + 0.1% BSA) and goat anti-rabbit IRDye 680 (Licor, 1:10,000 in TBST + 5% filtered milk) for 3 hours at room temperature. Loading was checked by staining with Revert total protein stain (Licor). Q1r is the Med15 fragment encompassing aa 120-277. 12PQ, 12L, NAB, HsRev, 24PQ and are substitutions of the 12Q region within Q1r consisting of the sequences shown in Fig. 3A. 0Q is a deletion of the 12Q from the Q1r construct. K is the Med15 fragment encompassing the KIX domain, aa 1-277. **(B)** N-Ubiquitin-Med15 protein levels per construct after normalization to total protein as determined with REVERT total protein stain (Licor) and made relative to Q1R which was set to 100%.

**Table S1. Strains**

| Strain | Relevant Genotype | Parental Strain &/or Plasmids | Source or Reference |
|---|---|---|---|
| BY4742 | MATα *his3Δ1 leu2Δ0 lys2Δ0 ura3Δ0* | (OY235) | [2] |
| (OY320) | *med15Δ0*::KanMX4 | BY4742 deletion collection | [2] |
| JF1368 | MATa *med15Δ*::*TRP1 ura3-52 lys2-801 ade2-101 his3Δ200 leu2Δ1* | | Fassler Lab Collection |
| CM67 | MATa *med15Δ0 his3Δ1 leu2Δ0 ura3Δ0 GAL11*-D7P | (OY383) BY4741 | [3] |
| CM66 | MATa *med15Δ0 his3Δ1 leu2Δ0 ura3Δ0 GAL11*-D30P | (OY384) BY4741 | [3] |
| MaV103 | *leu2-3,112 trp1-901 his3Δ200 ura3-52 ade2-101 gal4Δ gal80Δ* $cyh2^R$ $can1^R$ *GAL1::HIS3@LYS2 GAL1::lacZ SPAL10::URA3@ura3* | (OY219) | [4] |
| BY4716 | MATα *lys2Δ0* | (OY195) | [2] |
| JF2624 | MATα *leu2-3,112 trp1Δ1 ade2-101 GAL1::lacZ GAL1::HIS3* | Haploid from a cross between MaV103 and BY4716 | This study |
| JF2626 | MATa *leu2-3,112 trp1Δ1 lys2Δ0 gal80Δ GAL1::lacZ GAL1::HIS3* | Haploid from a cross between MaV103 and BY4716 | This study |
| JF2629 | MATα *med15Δ0*::KanMX4 *leu2-3,112 trp1Δ1 ade2-101 GAL1::lacZ GAL1::HIS3* | JF2624 | This study |
| JF2631 | MATa *med15Δ0*::KanMX4 *leu2-3,112 trp1Δ1 lys2Δ0 gal80Δ GAL1::lacZ GAL1::HIS3* | JF2626 | This study |

**Parenthetical names (OY###) are lab-specific storage designations.**

**Table S2. Plasmids**

| Plasmid | Details | Other Comments | Source or Reference |
|---|---|---|---|
| pRS315 | *LEU2*, CEN | (SV31) | [5] |
| pRS313 | *HIS3*, CEN | (SV32) | [5] |
| pRS416 | *URA3*, CEN | (SV23) | [5] |
| pRS315 M-WT | Mini-*MED15;* the internal region of Med15 (aa 116 to 277) plus the Mediator association domain (aa 799 to 1081) located between the promoter and terminator regions of S288C *MED15* in pRS315 (*LEU2*, CEN) | (SV286) | [6] |
| pLGZ-2LexA | Episomal LacZ reporter (*URA3*, 2μ) | (SV287) | [6] |
| pCL313/TIC | (pRS313 derived) copper inducible hGRτ1-LexA (*HIS3*, CEN) | (SV288) | [6] |
| pDC2141 | BY4742 *MED15* in pRS315 (*LEU2*, CEN) | SV286 gap repair | This study |
| pJF787 | S288C *MED15* locus in pRS416 (*URA3*, CEN) | | Fassler Lab Collection |
| pDC2285 | BY4742 153-687Δ *MED15* in pRS315 (*LEU2*, CEN) | | This study |
| pDC2286 | BY4742 46-619Δ *MED15* in pRS315 (*LEU2*, CEN) | | This study |
| | PARTIAL *MED15* GENE FRAGMENTS | | |
| (GC1012) | Topo 2.1 with Q2:*Afe*I gBlock | | This study |
| (GC1013) | Topo 2.1 with Q3:*Bmg*BI gBlock | | This study |
| (GC1014) | Topo 2.1 with Q2:*Afe*I Q3:*Bmg*BI gBlock | | This study |
| (GC1025) | Topo 2.1 with KIX gBlock | | This study |
| pDC2178 | Topo 2.1 with Q1:AfeI Q1R region from Mini-*MED15* (aa 116-277) | | This study |
| pDC2185 | 1x glycine spacer ligated into AfeI digested pDC2178 | | This study |
| pDC2253 | 6Q ligated into AfeI digested pJF2178 | | This study |
| pDC2255 | 12PQ sequence ligated into AfeI digested pDC2178 | | This study |
| pDC2256 | 24PQ- sequence ligated into AfeI digested pDC2178 | | This study |
| pDC2297 | LQ sequence ligated into AfeI digested pDC2178 | | This study |
| pDC2298 | FrHs sequence ligated into AfeI digested pDC2178 | | This study |

| | | | |
|---|---|---|---|
| **pDC2299** | **RvHs sequence ligated into AfeI digested pDC2178** | | **This study** |
| **pDC2300** | **NAB sequence ligated into AfeI digested pDC2178** | | **This study** |
| | **KIXQ2Q3Δ-*MED15* Q1 VARIANTS** | | |
| **pDC2136** | **KIXΔ (0:0:0) SV286 digested with *Pst*I and gap repaired with 0-Q1 PCR and Q2:*Afe*I Q3:*Bmg*BI gBlock** | | **This study** |
| **pDC2138** | **KIXΔ (12:0:0) as above with 12-Q1 PCR** | | **This study** |
| **pDC2144** | **KIXΔ (24:0:0) as above with 24-Q1 PCR** | | **This study** |
| **pDC2146** | **KIXΔ (36:0:0) as above with 36-Q1 PCR** | | **This study** |
| **pDC2147** | **KIXΔ (47:0:0) as above with 47-Q1 PCR** | | **This study** |
| **pDC2149** | **KIXΔ (12:0:0) as above with 12-Q1 PCR** | | **This study** |
| **pDC2150** | **KIXΔ (19:0:0) as above with 19-Q1 PCR** | | **This study** |
| **pDC2151** | **KIXΔ (21:0:0) as above with 21-Q1 PCR** | | **This study** |
| **pDC2152** | **KIXΔ (22:0:0) as above with 22-Q1 PCR** | | **This study** |
| **pDC2257** | **KIXΔ (1xG-rich:0:0) pDC2136 partially digested with AfeI and gap repaired with Q1R PCR from pDC2185** | | **This study** |
| **pDC2260** | **KIXΔ (6:0:0) as above with Q1R PCR from pDC2253** | | **This study** |
| **pDC2262** | **KIXΔ (1xPQ:0:0) as above with Q1R PCR from pDC2255** | | **This study** |
| **pDC2263** | **KIXΔ (2xPQ:0:0) as above with Q1R PCR from pDC2256** | | **This study** |
| **pDC2293** | **KIXΔ (12L:0:0) as above with Q1R PCR from pDC2293** | | **This study** |
| **pDC2291** | **KIXΔ (15LQ:0:0) as above with Q1R PCR from pDC2297** | | **This study** |
| **pDC2294** | **KIXΔ (FrHs:0:0) as above with Q1R PCR from pDC2298** | | **This study** |
| **pDC2295** | **KIXΔ (RvHs:0:0) as above with Q1R PCR from pDC2299** | | **This study** |
| **pDC2292** | **KIXΔ (NAB:0:0) as above with Q1R PCR from pDC2300** | | **This study** |
| **pDC2217** | **KIXΔ-*MED15*** | | **This study** |
| | **SPLIT UBIQUITIN** | | |
| **(GC408)** | **Empty CUb-RURA3** | | **addgene 131163** |
| **pDC2279** | **$MSN2_{TAD}$ CUb-RURA3** | | **This study** |

| | | | |
|---|---|---|---|
| **pDC2278** | **Empty Nub with N terminal CUP1 promoter and 3' UTR from MED15 (see Methods for details) in pRS315** | | **Built from addgene 131169** |
| **pDC2283** | **KIX Nub: KIX fragment gap repaired into PstI digested pDC2279** | | **This study** |
| **pDC2282** | **KQ Nub: KQ fragment gap repaired into PstI digested pDC2279** | | **This study** |
| **pDC2284** | **Q1R Nub: Q1R fragment gap repaired into PstI digested pDC2279** | **(12Q)** | **This study** |
| **pDC2301** | **0Q-Q1R Nub: 0Q-Q1R fragment gap repaired into PstI digested pDC2279** | | **This study** |
| **pDC2304** | **6Q-Q1R Nub: 6Q-Q1R fragment gap repaired into PstI digested pDC2279** | | **This study** |
| **pDC2302** | **24Q-Q1R Nub: 24Q-Q1R fragment gap repaired into PstI digested pDC2279** | | **This study** |
| **pDC2303** | **Spacer-Q1R Nub: Spacer-Q1R fragment gap repaired into PstI digested pDC2279** | | **This study** |
| **pDC2305** | **12PQ-Q1R Nub: 12PQ-Q1R fragment gap repaired into PstI digested pDC2279** | | **This study** |
| **pDC2306** | **24PQ-Q1R Nub: 24PQ-Q1R fragment gap repaired into PstI digested pDC2279** | | **This study** |
| **pDC2307** | **12L-Q1R Nub: 12L-Q1R fragment gap repaired into PstI digested pDC2279** | | **This study** |
| **pDC2280** | **NAB3-Q1R Nub: NAB3-Q1R fragment gap repaired into PstI digested pDC2279** | | **This study** |
| **pDC2281** | **RvHs-Q1R Nub: RvHs-Q1R fragment gap repaired into PstI digested pDC2279** | | **This study** |

**Parenthetical names (SV### and GC###) are lab storage designations.**

**Table S3. Primers**

| Primer | Sequence (5’->3’) | Use |
|---|---|---|
| ***MED15* F-245** | **GGATGAGGATGATGAAGGTGC** | **Screen & Amplify *MED15* ORF** |
| ***MED15* F+334** | **GCAACAGCGCCAATAATATGAATG TCAAT** | **Q1 PCR, Amplify Central Q-rich Region & Sequencing** |
| ***MED15* R+529** | **GTGCCACTTTCATCTGGTTC** | **Q1 PCR, Screen & Sequencing** |
| **MED15 F+854** | **CACAAAATACCGTACCAAACGTCC** | **Q2 PCR** |
| ***MED15* F+1204** | **GCCGCTTTCTCTCAGCAACAGAAC** | **Screen** |
| ***MED15* F+1682** | **CCCCACAAGTCTACATCATCACAA AG** | **Q3 PCR** |
| ***MED15* R+1781** | **CCCTCGGCACATTTTTCTAGGATC** | **Q2 PCR and Sequencing** |
| ***MED15* F+1924** | **GGCACTACTTCTACTGGAAAC** | **Screen** |
| ***MED15* R+2143** | **CATTAGCCATTGCCGAATAATTAG CCACACCAGG** | **Screen & Amplify Central Q-rich Region** |
| ***MED15* R+2385** | **CCTAGGAGTGGGTTGTTGACTAG** | **Q3 PCR and Sequencing** |
| ***MED15* F+2693** | **AGATGCTTACGTCATGCACTATCC** | **Sequencing** |
| ***MED15* R+2990** | **GGGTTGCCGACATCCATATTAG** | **Sequencing** |
| ***MED15* R+3498** | **GTACTGATGATAGTCAAGTCCATT G** | **Screen & Amplify *MED15* ORF** |
| **M13 F** | **GTAAAACGACGGCCAG** | **Screen & Sequencing** |
| **M13 R** | **CAGGAAACAGCTATGAC** | **Screen & Sequencing** |
| **Q1R gap repair F** | **GATCGCATCGAATTCCTGCAGCCC GGGGGATCCAATATGAATGTCAAT ATGAATCTAAAC** | **Q1R PCR** |
| **Q1R gap repair R** | **GATTTGTATTTGGGGTTGACTTAG CGGTATTCTGCAGCAAAGGGTTGT TGTTCGCACTTG** | **Q1R PCR** |
| **0Q F** | **CAGGTTGCGCAATTAAGAAATAGC GCTAGGCGTCAATTGACTCCT** | **Amplify Q1-0 *Afe*I** |
| **0Q R** | **AGGAGTCAATTGACGCCTAGCGCT ATTTCTTAATTGCGCAACCTG** | **Amplify Q1-0 *Afe*I** |
| **KIX gBlock F** | **TATAGTGCCGTACTCAAAGATC** | **Amplify KIX gBlock** |
| **KIX gBlock R** | **TGTTGCGCAACCTGTTGC** | **Amplify KIX gBlock** |

**Table 4 qRT-PCR Primers**

| Gene | Product Size | Efficiency | Forward Primer Sequence 5'->3' | Reverse Primer Sequence 5'->3' |
|---|---|---|---|---|
| *AHP1* | 179 | 1.84 | GGAAGTTGACCAAGTGATCG | CTACCACTCCAGTAAACACC |
| *ALG9* | 142 | 2.02 | GGATAGTGGCTTTGGTGAAC | CAGGAAAGAACTTGGGAAGTG |
| *ARG3* | 126 | 2.22 | GATGGCATGGATTGGTGATG | TTCATCGACAATATCGGAATCC |
| *ARG5/6* | 183 | NT* | CTGTGAACCTTTCGTTGAGAC | CGAACCAGTAGCATAACAACC |
| *GLK1* | 133 | 1.97 | AACACGGACTCAATGACGTC | AACTTGTCACGCAACTCCTG |
| *HSP12* | 161 | 2.03 | CATACGCTGAACAAGGTAAGG | CTTGGTCTGCCAAAGATTCAC |
| *MED15* | 297 | 2.02 | AGATGCTTACGTCATGCACTATCC | GGGTTGCCGACATCCATATTAG |
| *MET10* | 109 | NT | ACTCAATTGCCTCCTCTCAG | CTTAGAAGCTTGACCGTACC |
| *SSA1* | 167 | 1.97 | CCAACAAAGAAGTCCGAGATC | GACTTCAATTTGTGGGACACC |
| *TFS1* | 167 | 1.98 | CCCAATTCCAGTTCACGTTC | AGTCGCATTCTACCAAGTGG |

### References


1. Gruber, M., J. Soding, and A.N. Lupas, *Comparative analysis of coiled-coil prediction methods.* J Struct Biol, 2006. **155**(2): p. 140-5.
2. Brachmann, C.B., et al., *Designer deletion strains derived from Saccharomyces cerevisiae S288C: a useful set of strains and plasmids for PCR-mediated gene disruption and other applications.* Yeast, 1998. **14**(2): p. 115-32.
3. Miller, C., et al., *Mediator phosphorylation prevents stress response transcription during non-stress conditions.* J Biol Chem, 2012. **287**(53): p. 44017-26.
4. Vidal, M., et al., *Reverse two-hybrid and one-hybrid systems to detect dissociation of protein-protein and DNA-protein interactions.* Proc Natl Acad Sci U S A, 1996. **93**(19): p. 10315-20.
5. Sikorski, R.S. and P. Hieter, *A system of shuttle vectors and yeast host strains designed for efficient manipulation of DNA in Saccharomyces cerevisiae.* Genetics, 1989. **122**(1): p. 19-27.
6. Kim, D.H., et al., *Functional conservation of the glutamine-rich domains of yeast Gal11 and human SRC-1 in the transactivation of glucocorticoid receptor Tau 1 in Saccharomyces cerevisiae.* Mol Cell Biol, 2008. **28**(3): p. 913-25.
